# To hear or not to hear: How Selective Tidal Stream Transport Interferes with the Detectability of Migrating Silver Eels in a Tidal River

**DOI:** 10.1101/2023.07.13.548823

**Authors:** Benedikt Merk, Leander Höhne, Marko Freese, Lasse Marohn, Reinhold Hanel, Jan-Dag Pohlmann

## Abstract

Acoustic telemetry provides valuable insights into behavioural patterns of aquatic animals such as downstream migrating European eels (*Anguilla anguilla*). The behaviour of silver eels during the migration is known to be influenced by environmental factors, yet so is the performance of acoustic telemetry networks. This study focusses on quantifying the impact of these environmental factors on both, migration behaviour and receiver performance, in order to determine possible limiting conditions for detecting tagged eels in tidal riverine areas and estuaries. A dominance analysis of the selected models describing migration speed, activity and receiver performance was conducted following 234 silver eels that were tagged with acoustic transmitters and observed by a receiver network in the Ems River during two subsequent migration seasons. The results suggest a passive locomotion of silver eels during their downstream migration by taking advantage of selective tidal stream transport (STST) It is further shown that water temperature, salinity, turbidity, precipitation, and especially current velocity were major parameters influencing migration activity and speed. At the same time, analyses of the detection probability of tagged eels under varying environmental conditions indicated a decreased receiver performance during high current velocities, resulting in a coincidence of high migration activity and reduced detection probability. Correspondingly, there is a risk that particularly during phases of increased activity, due to limited telemetry performance, not all fish will be detected, resulting in an underestimation of migration activity. To avoid misleading interpretations and underestimates of migration numbers of eels and other migratory fish using STST, this study highlights the need to conduct range tests and adjust the receiver placement in areas and conditions of high current velocities.

## Introduction

With increasingly sophisticated technology, advanced acoustic telemetry networks allow for detailed behavioural analyses of target organisms from cubozoans (Gordon & Seymour, 2009) to whale sharks (Cagua et al., 2015). This passive methodology is based on acoustic signal transmission between installed receivers, forming a receiver network, and transmitter tags, which are implanted or externally attached to a target organism. Many behaviour-related research questions including movement patterns, migration speed and timing, predation events and 3-dimensional spatial distribution can be investigated with this methodology causing relatively little impact on the organism’s life (Béguer-Pon et al., 2012; Melnychuk, 2012; Kessel et al., 2014; Hussey et al., 2015; Béguer-Pon et al., 2018a). For a detailed assessment and prediction of animal behaviour, species distribution and migratory patterns, especially in light of conservation efforts, a combination of acoustic telemetry data with abiotic environmental factors can be of great benefit (Matley et al., 2022).

The behaviour of migratory fish is known to be largely influenced by prevailing environmental conditions. In general, it is hypothesised that fish prefer to migrate during certain environmental conditions to save energy, maximize survival and avoid predation by other fish, birds and marine mammals (Sandlund et al., 2017). In the case of the critically endangered European Eel (*Anguilla anguilla*), high current velocities (Trancart et al., 2018) and river discharge are linked with increased migration activity (Vøllestad et al., 1986; Cullen & McCarthy, 2003; Bultel et al., 2014; Marohn et al., 2014; Reckordt et al., 2014) and migration speed (Vøllestad et al., 1986; Stein et al., 2016). It is hypothesised that selective tidal stream transport (STST) plays an important role in the riverine and coastal migration, allowing for energy conservation of the migrating species by ascending into the water column, when the tidal flow equals the migratory direction and settling on the bottom or sheltered areas during opposite tide intervals (Walker et al., 1978; Forward and Tankersley 2001; Verhelst et al., 2018). The usage of STST has been proven for various fish species, including the silver eel stage of the American eel (*Anguilla rostrata*) (Béguer-Pon et al., 2014), glass eel stage of several *Anguilla* species (Sugeha et al., 2001; Trancart et al., 2012), Green sturgeon (*Acipenser medirostris*) (Kelly et al., 2020), Thinlip grey mullet (*Chelon ramada*) (Trancart et al., 2012) and flounder larvae (*Platichthys flesus*) (Jager et al. 1999; Trancart et al., 2012). For the European silver eels, studies by Barry et al. (2016), Huisman et al. (2016) and Verhelst et al. (2018) recorded increased migration activity during ebb tide, strongly suggesting the usage of ebb-tide transport during their downstream migration, while a study by Bultel et al. (2014) could not confirm this observation. Additionally, meteorological effects like periods with increased precipitation, often coinciding with increased water level and flow (Stein et al., 2016; Trancart et al., 2018) as well as lowered atmospheric pressure (Cullen & McCarthy, 2003) and wind (Cullen & McCarthy, 2003; Reckordt et al., 2014) are mentioned as possible triggers of migration. Besides, water temperature also influences the migration activity of European eels (Vøllestad et al., 1986; Lobón-Cerviá & Carrascal, 1992; Bruijs & Durif, 2009), as their ectothermic metabolism shows limited activity during high (linked to decreasing oxygen concentrations or other mechanisms) or low temperatures (Walsh et al., 1983; Claësson et al., 2016). According to Lennox et al. (2018), eels avoid migrations during daylight and are thought to migrate preferentially during new moon, hence reducing their vulnerability to predators (Barry et al., 2016; Stein et al., 2016). However, the role of the lunar cycle could not be corroborated by other studies (Marohn et al., 2014; Reckordt et al., 2014), as it can be inhibited by other conditions, such as increased turbidity (Cullen & McCarthy, 2003; Durif et al., 2003, Höhne et al., under review).

In case of the critically endangered European eel, knowledge about site-specific environmental drivers of migration can also be used to predict migration events and implement protection measures, e.g. the shutdown of hydropower turbines (Teichert et al., 2020) or the determination of closed seasons in the absence of actual monitoring data. However, apart from testing environmental influences on migratory fish behaviour, an important and often underestimated prerequisite for the use of telemetry is to estimate the detection probability of the installed telemetry network during different environmental conditions and hence the validity of the respective data.

Hydrological- and meteorological parameters affect the underwater acoustic landscape, with implications for the transmission of sounds. Therefore, these factors can limit the detection of emitted acoustic telemetry signals drastically, inducing uncertainties and possibly even erroneous conclusions of the main biological research subject (Kessel et al., 2014). Past studies determined current velocity and turbidity as main influencing factors in coastal systems (Mathies et al., 2014; Reubens et al., 2019). Additionally, waves and wind can induce under water noise, possibly impairing successful signal transmission. In general, the presence of air bubbles (e.g. entrapped by precipitation, waves and currents) and sediment particles can cause an undirected scattering or absorption of the sound waves resulting in a higher signal loss (Stocks et al., 2014; Reubens et al., 2019). Moreover, ambient noise, originating from anthropogenic sources, e.g., nautical traffic, rattling of buoy chains, is known to potentially conceal transmitter pings (Kessel et al., 2014; Mathies et al., 2014; Reubens et al., 2019). Other environmental factors, potentially interfering with transmitter-receiver interaction include increased salinity and water temperature favour signal transmission (Yuwono et al., 2014), by changing the physical properties (e.g., density and viscosity) and allowing for an improved sound propagation (Makar 2022). Beside the environmental impacts, also technical parameters such as receiver tilt angle and position of the receiver’s hydrophone in relation to the signal-emitting tag are important factors for the performance of the receiver network (Reubens et al., 2019). Additionally, the detection probability scales with the power output of the used tags (Stott et al., 2021). A sentinel tag approach with fixed tags or receiver internal tags, communicating with each other in a chain of devices, enables a continuous control of the detection probability at fixed distances (Reubens et al., 2019).

This paper analyses the environmental drivers of the downstream migration of silver eels, together with their influences on detection probability within an acoustic telemetry network installed in the tidal- and estuarine area of the German River Ems. The aim was to determine the relative importance of factors in explaining migration activity, -speed and detection range, which collectively determine the probability of successful detection by acoustic telemetry. This allowed for the identification of possible conditions that favour increased and faster migrations, while simultaneously lowering the effective range of the hydrophone network, causing a possible bias in the data analysis during main migration events.

## Methods

### Study Area

In order to study aspects of migration behaviour and detectability of tagged European eels across two migration seasons, 29 telemetry receivers (Model VR2Tx, Vemco Ltd Halifax, Canada), forming seven acoustic arrays, were set up along the tidal and inland area of the River Ems. The entire study area comprised 109.3 km of the Ems River between the town of Meppen and the seaward boundary of the Dollart estuary, which forms the German-Dutch border. In this study the focus was set on the tidal area stretching from the tidal weir in Herbrum to the Dollart Estuary including receiver arrays 4 to 6 (Figure 1). This highly anthropised part of the Ems River is characterised by a straightened and deepened river bed as well as a muddy bottom structure, high sediment loads and regular tidal cycles. Receiver Array 7 consists of a chain of 12 telemetry receivers covering the width of the Dollart Estuary, forming the direct connection and entry into the North Sea.

**Figure 1:**
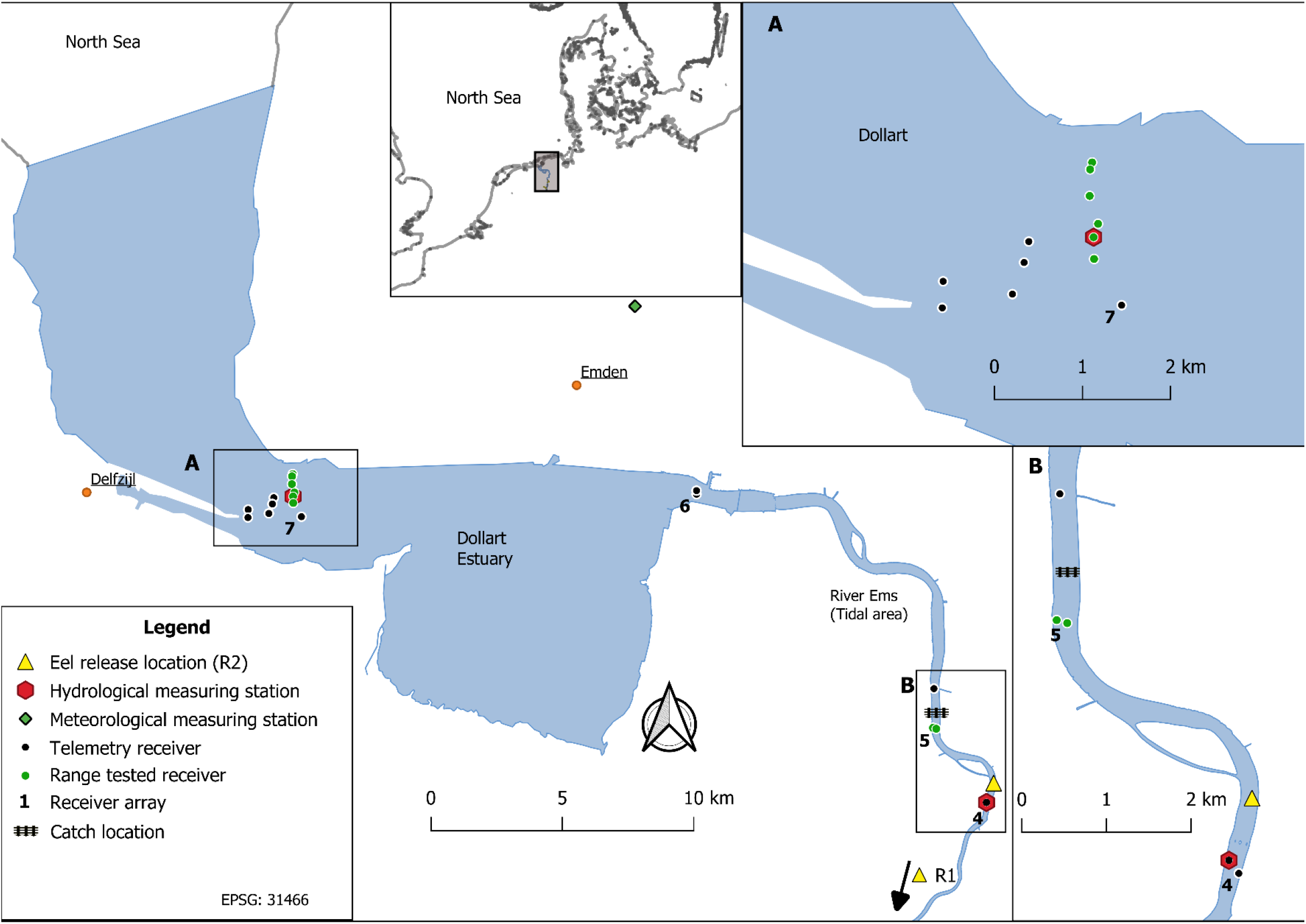
Study area including the telemetry network along the Ems River, with detailed maps of the range-tested receiver locations (A-Estuary, B-Tidal area). Catch and Release locations, nvironmental measuring stations and receiver arrays (4-7) are also marked

### Telemetry Network and Range Tests

The telemetry network consisted of 19 acoustic receivers (Vemco VR2Tx) with built-in tags emitting and receiving signals with a frequency of 69 kHz (Vemco, 2016) in the tidal area and estuary of the Ems River. Receivers were attached to anchor chains of navigation buoys (hereafter referred to as buoy chains) at a water depth of 2-3 m with the exception of two receiver units placed in shallow areas that necessitated a custom mooring system with a concrete anchor, polyester rope, and a floatation buoy. The assessment of the detection probability during different conditions was synchronised with the downstream migration of tagged silver eels in order to directly assess the effect of environmental conditions on their behaviour.

The detection probability is the relation between emitted and detected acoustic signals within telemetry receiver arrays. In order to test the acoustic detection probability and range of the receivers in the tidal area (Array 5) and estuary (Array 7) constantly over a longer time period, a sentinel approach, with fixed positions of signal emitters and receivers was used. The focal receiver arrays were located in vicinity of stationary hydrological measuring stations, maximizing representativity of the environmental data. The arrays were unobstructed by structures such as curves and shore line irregularities, which would limit the signal transmission range or fully disrupt transmission (Walsh et al., 2012). Selected receivers (see green markings in Figure 1) and range test tags were set to emit acoustic signals (“pings”) every 10 minutes. Within the estuarine receiver array (Figure 1A), the detection probability was tested for six receivers over different distances between 85 and 1097 m (Appendix Table A1). In the tidal array, the detection probability was measured using a receiver pair positioned 129 m apart.

Prior to the continuous assessment of detection probability, initial boat drifts were conducted in the tidal area (setup see Appendix Text) from which the 50%-detection-range (i.e., the range at which 50% of all pings are detected) was determined. This method helps to standardise the relation of detected and non-detected pings and to understand how this relation changes during changing environmental conditions (Brownscombe et al., 2020). The receiver pair in the tidal area (Array 5) was chosen to best represent the calculated 50%-detection-range (corresponding to the 53%-detection range instead of the 50%-range).

The ping intensity of the selected receivers in the tidal and estuarine array was set to 142 dB to avoid interference of ping originating from implanted V9-tags pinging at 146 dB. The receivers were maintained regularly, in order to change batteries and avoid accumulation of biofilm, sessile organisms and debris on the hydrophone.

Due to difference in the environmental conditions and technical setup the following analysis were conducted separately for the estuarine (Array 6-7) and tidal area (Array 3-6) (Figure 1).

### Sampling

In line with this study, 234 female silver eels were tagged with acoustic telemetry tags (Vemco V9, 9 mm diameter; 2.0 g in water, min. 409 days of battery life), 35 of which were classified as Stage FIV and 199 as stage FV based on Durif et al. (2005, 2009). The transmission rate was set at 60 seconds with a random variation of ±20 seconds, to avoid signal overlaps of multiple tags (Vemco, 2020). All eels were caught in a fixed stow net, located in the tidal area within array 5. The net was emptied daily by a local fisher and eels were stored in a holding box and tagged within two days after catch. The tagging process was initiated by anaesthetization with clove oil (Walsh and Pease, 2002). Once the eels reached the stage of narcotic immobility, total length, weight, eye diameter and pectoral fin length were measured, and an acoustic transmitter was surgically implanted into the body cavity. The incision was closed with two stitches, using a slowly absorbable monofilament suture (Surgicryl monofilament DS 24, 3.0 (2/0), SMI AG, St. Vith, Belgium) (Thorstad et al., 2013).

Additionally, a T-bar anchor tag was inserted in the epaxial musculature. Tagged eels were allowed to recover for 1-8 hours in a dark tank with air dispersal to recover and released at an inland (R1) or tidal release location (R2). During the transportation to the R1 location, freshwater from an inland river site was added to the recovery tank in order to enable a gradual acclimatisation to lower salinity levels. Prior to release it was ensured that eels regained active swimming and flight reflexes. Since eels released in the inland- or tidal region did not differ in survival (Höhne et al., under review) and migration speed (p=0.986, Figure Appendix 1), both groups were retained in the analyses.

The observation period covered two main migration seasons, defined as the period from 15.10.2020 – 27.02.2021 and 29.09.2021 – 27.02.2022 (Höhne et al., under review). In the first season the observation time was limited by drift ice, necessitating temporal removal of the stow nets and some receivers, thus no data was generated after the 05.02.2021.

### Environmental Data

Hydrological parameters (water temperature, current speed, current direction, water level, electric conductivity, salinity, turbidity, O2-conc.) were measured and obtained by measuring stations in the Dollart estuary (within Array 7) and in the tidal area (close to Array 5), operated by the local water authority WSV Ems-Nordsee (Figure 1). Meteorological data (wind speed, precipitation) originating from the official German weather Agency Deutscher Wetterdienst (DWD) in Emden and Moormerland were associated with the estuary and tidal area, respectively as these were the closest measuring stations. Further, illuminated moon fraction was obtained from the R package “suncalc” for the geographic position 52.23196 N, 7.412325 E, located in the centre of the study area. Data was binned hourly based on the parameter with the lowest temporal resolution. Current directions between 31° to 211° in the estuary and 106° to 286° in the tidal region were defined as upstream currents and marked with a negative prefix for the analysis of the eel behaviour. Additionally, accumulated precipitation of the last seven days was computed by summing up the hourly precipitation values. Moreover, the water level difference was calculated by subtracting water level at hour *n* - 1 from water level at hour *n*. Although environmental influences were the main interest of this study, receiver internal data (tilt angle and surrounding noise) and the azimuth, following the definition of Reubens et al. (2019), were included as control variables for the analysis of the detection probability. A low azimuth angle indicated that the receiver hydrophones are facing toward each other, while a high angle implied opposite facing directions, hence reducing the detection probability, as the receiver body overshadows the hydrophone. For the tidal area, hydrological data were missing from 03.- 16.12.2021, as the measuring instruments were not operational.

### Statistical analysis

Statistical analyses aimed at identifying the effect of environmental conditions on silver eel migration activity and speed, as well as acoustic detection probability. Analyses were conducted separately for the tidal (Figure 1B) and estuarine region (Figure 1A). Accordingly, six generalised linear models were computed, each starting with a maximum model as described in Table 1.

**Table 1:** List of all computed statistical models, with model type, dependent and independent variables. Variables in the column “excluded correlates” were eliminated from the maximum model due to collinearity with other parameters. GLM- Generalised linear model, * - interaction terms.

| Abbreviation | Model (Family) | Maximum Model | Excluded Correlates |
| --- | --- | --- | --- |
| MSE (Estuary) | GLM (Gamma, log-link) | Migration speed ~ Wind speed + Precipitation + Acc. Precipitation + Water Temperature + Current velocity + Salinity + Moon fraction + Water level diff. | O<sub>2</sub>-Conc.<br>El. Conductivity<br>Water level diff. |
| MST (Tidal) | GLM (Gamma, log-link) | Migration speed ~ Wind speed + Precipitation + Acc. Precipitation + Turbidity + Water Temperature + Current velocity + Moon fraction + Water level diff. | O<sub>2</sub>-Conc.<br>Water level diff. |
| MPE (Estuary) | GLM (binomial) | Migration probability ~ Wind speed + Precipitation + Acc. Precipitation + Water Temperature + Current velocity + Salinity + Moon fraction + Last release | O<sub>2</sub>-Conc.<br>El. Conductivity<br>Water level diff. |
| MPT (Tidal) | GLM (binomial) | Migration probability ~ Wind speed + Precipitation + Acc. Precipitation + Turbidity + Water Temperature + Current velocity + Moon fraction + Last release | O<sub>2</sub>-Conc.<br>Water level diff. |
| DPE (Estuary) | GLM (binomial) | Detection probability ~ Wind speed + Precipitation + Acc. Precipitation + Water Temperature + Current velocity (no prefix) + Salinity + Water level diff. + Last release + Water level + Azimuth*Tilt + Distance*Noise | O<sub>2</sub>-Conc.<br>El. Conductivity |
| DPT (Tidal) | GLM (binomial) | Detection probability ~ Wind speed + Precipitation + Acc. Precipitation + Turbidity+ Water Temperature + Current velocity (no prefix) + Water level diff. + Last release + Water level + Azimuth*Tilt + Distance*Noise | O<sub>2</sub>-Conc.<br>El. Conductivity |

**Table 2:**
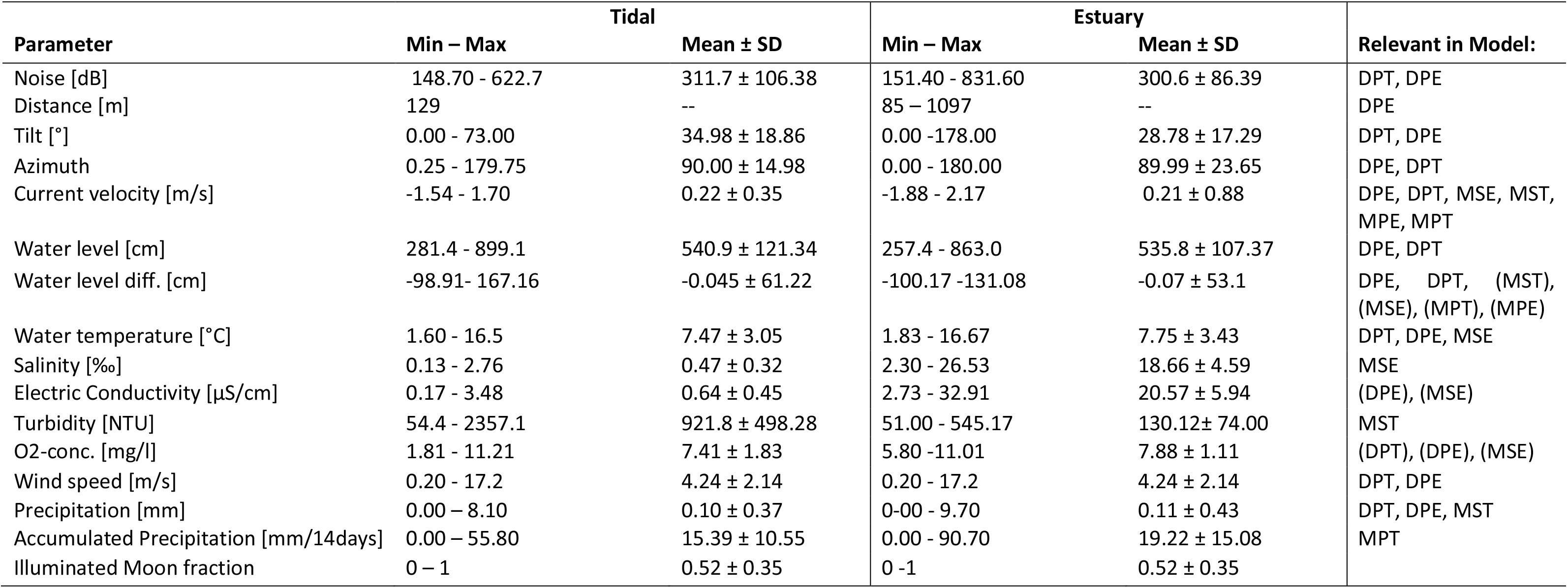
Summary of the surveyed environmental and technical parameters within the observation period, as well as their inclusion in the final models following model selection. Parentheses symbolize importance via a correlation. Model abbreviations: MS- Migration Speed, MP-Migration Probability, DP-Detection Probability, T-Tidal, E-Estuary

Environmental data were z-transformed (i.e., mean-centred and scaled to standard deviations) to facilitate immediate comparability within the observed range of values.

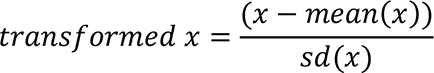

Collinearity was defined by a correlation coefficient over rho >0.7 (Spearman correlation-test). If two explanatory variables were correlated, only one of the two covariates was included into the maximum model.

The parameters of the maximum model were eliminated in a stepwise backward selection procedure (Zuur et al., 2009), based on AIC-values (Akaike Information Criterion, Akaike, 1987), to define the final minimum adequate model, prioritising models with fewer variables when the AIC was similar (AIC difference < 2). Selected models were validated by visual inspection of residuals over fitted values and normal-Q-Q plots.

Map creation, transect and distance calculation were conducted with QGIS (Vers. 3.14 “Pi “) (QGIS.org, 2021). All statistical analyses were performed in R version 4.1.0 (R Core Team, 2021) using packages “dominanceanalysis” (Vers.2.0.0) (Navarrete and Soares, 2020) and “parameters” (Vers.0.21.0) (Lüdecke et al., 2020) for the dominance analysis and packages “nlme” (Vers. 3.1-152) (Pinheiro et al., 2021) and “lme4” (Vers. 1.1-27.1) (Bates et al., 2015) for the initial GLMM- approach.

#### Analysis of migration speed

Migration speed was analysed in relation to prevailing environmental conditions. Downstream migration speed of an eel was calculated as the time difference (in h) between first detection on an upstream array and the first detection on a downstream array, divided by the distance between the two receivers in river-km. Only downstream movements that happened within 24 h (i.e., when an eel was detected at two arrays within 24 h) were retained in the analysis, to exclude non-directed downstream movements and to avoid unrepresentative means for environmental parameters over extended time periods. The environmental parameters were averaged over the duration of the downstream movement of an eel between two arrays. For the tidal area migrations between Array 3 and Array 6, and for the estuary migrations between Array 6 and 7 were considered. A generalised linear mixed-effects model with a random intercept for individual eel was fitted initially, but the random effect was subsequently excluded, as it had no impact on migration speed, increased comparability with the other models and the model was reduced to a GLM for the tidal- and estuarine analysis (model “MST” for tidal-, and “MSE” for estuarine area; Table 1).

To assess the active swimming of eels during the downstream migration, migration speed in relation to the current velocity (hereafter “relative migration speed”) was calculated, by dividing the migration speed by the current velocity, with the exclusion of moments during average stagnant water (<|0.01| m/s).

#### Analysis of migration activity

Migration activity was measured as the number of tagged silver eels exhibiting downstream movement, divided by the number of tagged individuals present in the observed area. Eels entering the study area through migrations from upstream or through release events increase the number of individuals in the area. Detections at Array 6 and 7 are defined as escaped from tidal area or estuary respectively and decrease the number of individuals present in the area. As no eel remaining in the study area within the first migration season, continued the migration in the second migratory period. Therefore, the number of individuals in the observed area was assumed to be 0 before the first release event of the second migration season at the 29.09.2021. This was done in order to minimize the effect of inactive (e.g., dead or remaining eels from the previous season) eels lowering the migration activity in the second observation period. The hourly binned environmental and technical factors were summarised for tidal cycles as the predominant biological cycle in the area. A tidal cycle is defined by period of continuous decrease or increase in water level. Therefore, the data set consisted of alternating ebb and flood tidal cycles. It was expected that many eels would continue their migration shortly after their release. To account for potential bias towards environmental conditions shortly after release events, typically followed by high migration activity, an independent variable (hours since last release event) was included in each model. The environmental influences on migration activity in the tidal area were tested with the MPT-model (glm, binomial family) and in the estuary with the MPE- model (glm, binomial family) (Table 1).

#### Analysis of the acoustic detection probability

To assess detection quality of the estuarine network array (Array 7), the relationship of distance (between two receiver units) and detection probability (i.e., the number of pings detected divided by the number of pings emitted at that given distance) was established. This was done on an hourly basis first and these datapoints were then summarised to get the distance detection probability relationship across the whole study period. From this relationship, the 50%- and 80%- detection-ranges (i.e., the distances at which 50% or 80% of emitted signals are detected) were calculated.

In the tidal area (Array 5), the given detection probability at 129 m (the distance of the observed receiver pair), was calculated (Brownscombe et al., 2020).

The environmental influences on receiver performance were tested with binomial GLMs using the detection probability as response variable for the estuarine analysis (model “DPE”) and for the tidal area (model “DPT”) (Table 1).

To calculate the chance of detection (CoD), the shortest transect a migrating eel could theoretically take in the estuary and tidal setup was determined with the intersect of the two overlapping detection radii of the receivers (Matthies et al., 2014). Transect length and migration speed of tagged eels (with average ping intervals of 60 sec) provided information about the possible emitted pings per transect (P/T) at a given detection probability (DP, e.g., 50% for estuarine, 53% for tidal area). This allowed to calculate the chance of detection (CoD) for these setups of a migrating eel with the following formula:

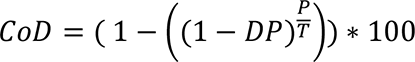

A high chance of detection validated the evaluation of the migratory patterns, as the number of undetected eels was rather low.

#### Dominance Analysis

The relative importance of a given environmental factor in a selected model was determined by a dominance analysis (Azen & Budescu, 2003; Azen & Traxel, 2009). Dominance analysis determines the relative importance of individual variables in a multivariate model based on each variable’s contribution to an overall model fit statistic (R²). The dominance analysis was based on Pseudo-Mc- Fadden-R² fit statistics (McFadden, 1974 and 1979) for binomial models (DPT, DPE, MPT, MPE) and Pseudo-Nagelkerke-R² values (Nagelkerke, 1991) for Gamma models (MSE, MST; Table 1).

Higher R² values reflect a better fit for the model and values between 0.2 and 0.4 for the Pseudo- McFadden-R² are considered to represent an excellent fit (McFadden, 1974 and 1979). The relative importance and effect direction were used to compare the parameteŕs influence on the three response variables in the tidal and estuarine analysis respectively. Thereby, parameters could be identified that affect both migration behaviour and detection probability simultaneously.

## Results

### Migration Speed

Average migration speeds of silver eels were 0.69 ± 0.41 m/s (mean ± SD), ranging from 0.05 to 1.44 m/s in the tidal area and 0.62 ± 0.44 m/s (mean ± SD) in the estuary, ranging from 0.18 to 1.86 m/s. Migration speed was faster during conditions of high current velocity, lower turbidity and high precipitation (n= 198, Pseudo-Nagelkerke-R²=0.545, model “MST”, Table 3). In the estuary, increased salinity, lower current velocities and water temperatures caused a slower migration (n= 142, Pseudo-Nagelkerke-R²=0.689, model ”MSE”, Table 3).

**Table 3:**
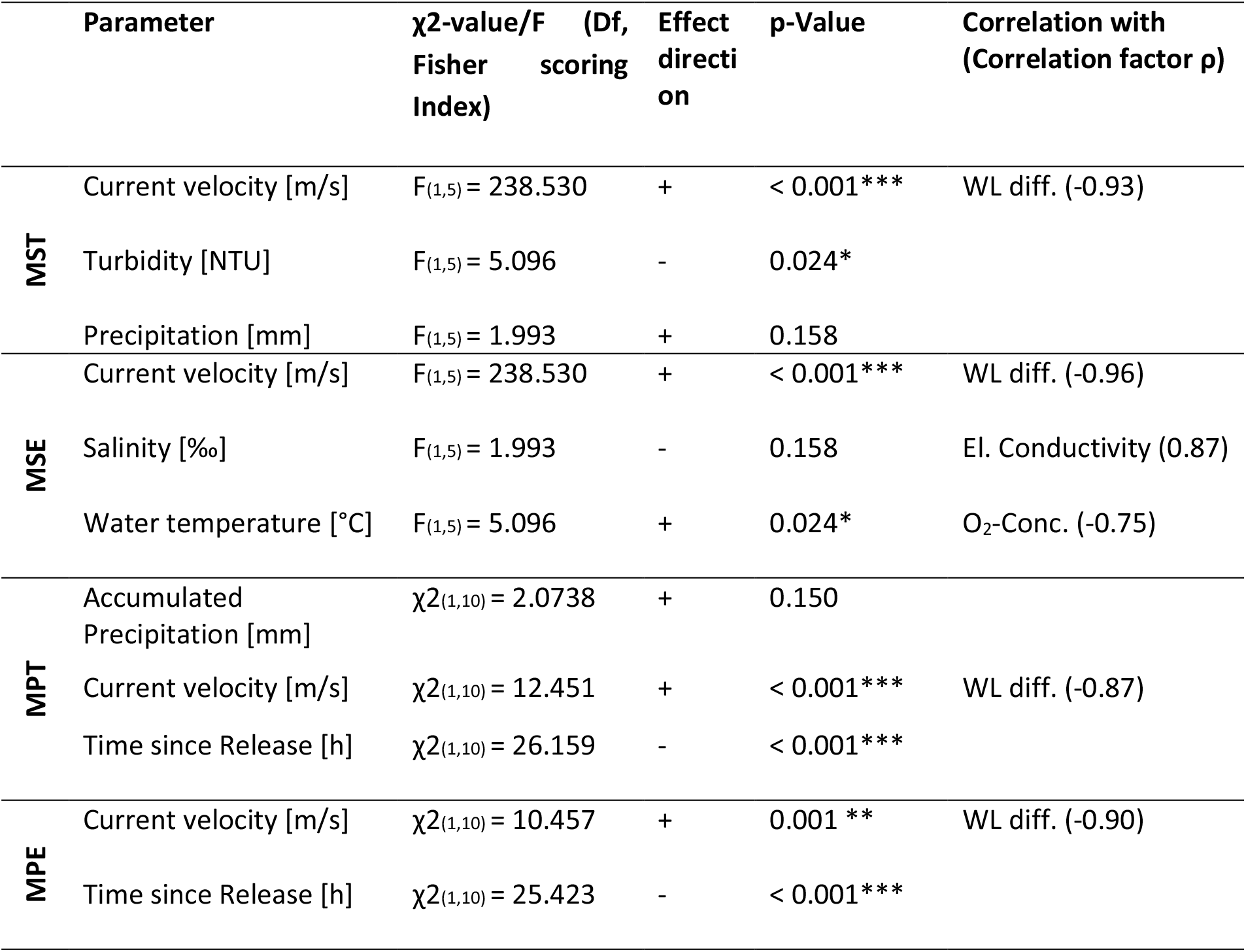
Summary of the minimum adequate models of factors influencing the migration speed and probability in the tidal area and estuary, Model abbreviations: MS- Migration Speed, MP-Migration Probability, DP-Detection Probability, T-Tidal, E-Estuary

The relative migration speed (swimming speed corrected for current velocity) was close to 1 in the tidal region and estuary, with exceptions during stagnant waters (Figure 2).

**Figure 2:**
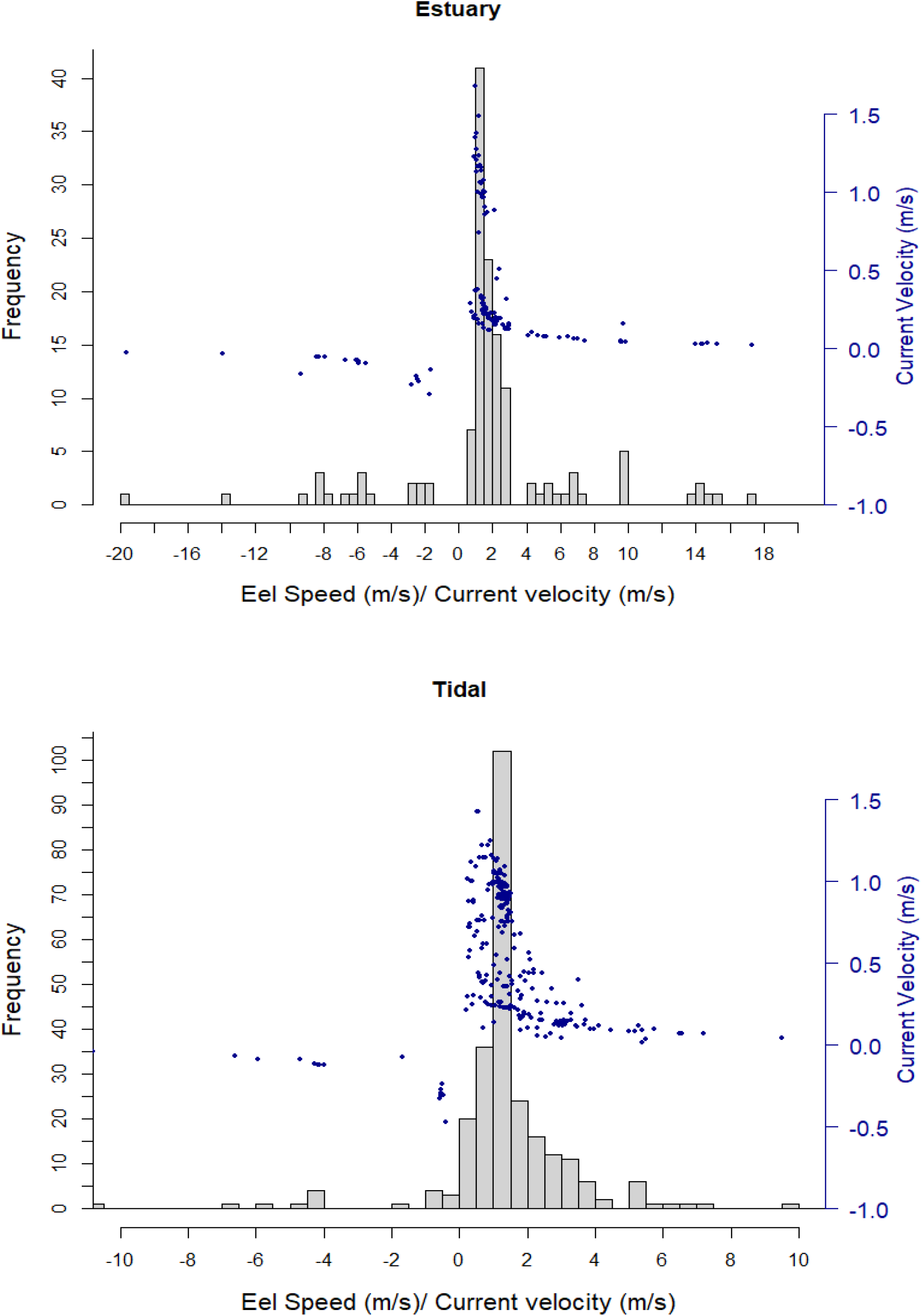
Histogram of the relative migration speed (Eel swimming speed/Current velocity) of eels in the estuary (top) and tidal area (bottom) with the blue dots marking the respective current velocity

### Migration activity

In total, 1080 tidal cycles within the observation period were analysed. 89.44 and 87.86% of the observed migration activity were recorded during ebb tides (i.e. tide cycles of downstream flow) for the tidal area and estuary, respectively. Migration probability increased with less time elapsed since a release event.

Besides that, accumulated precipitation and current velocity increased the migration activity in the tidal area (*N* = 323 downstream movements, Pseudo-McFadden-R²=0.24, model “MPT”, Table 3). In the estuary, high current velocities induced higher migration activity (*n=* 173 downstream movements, Pseudo-McFadden-R²=0.022, model “MPE”, Table 3).

### Receiver Performance

In the tidal area (Array 5), 12,682 pings were emitted by both receivers in the tidal system of which 6,803 pings were detected. The average detection probability at the 129 m distance was 53.64% (Figure 3). During conditions with an elevated water level, higher precipitation and higher azimuth the detection probability was increased while higher current velocity, noise, tilt, water level difference, wind and water temperature decreased the chance of receiving pings (Pseudo-McFadden-R²=0.489, model “DPT”, Table 4). In the estuary (Array 7) the six tested receivers emitted 219,714 pings, resulting in 144,168 detects by other receivers during the observation period. The 50%- detection range was evaluated at 197 m and the 80%- detection range at 98 m (Figure 4, Figure Appendix A2).

**Figure 3:**
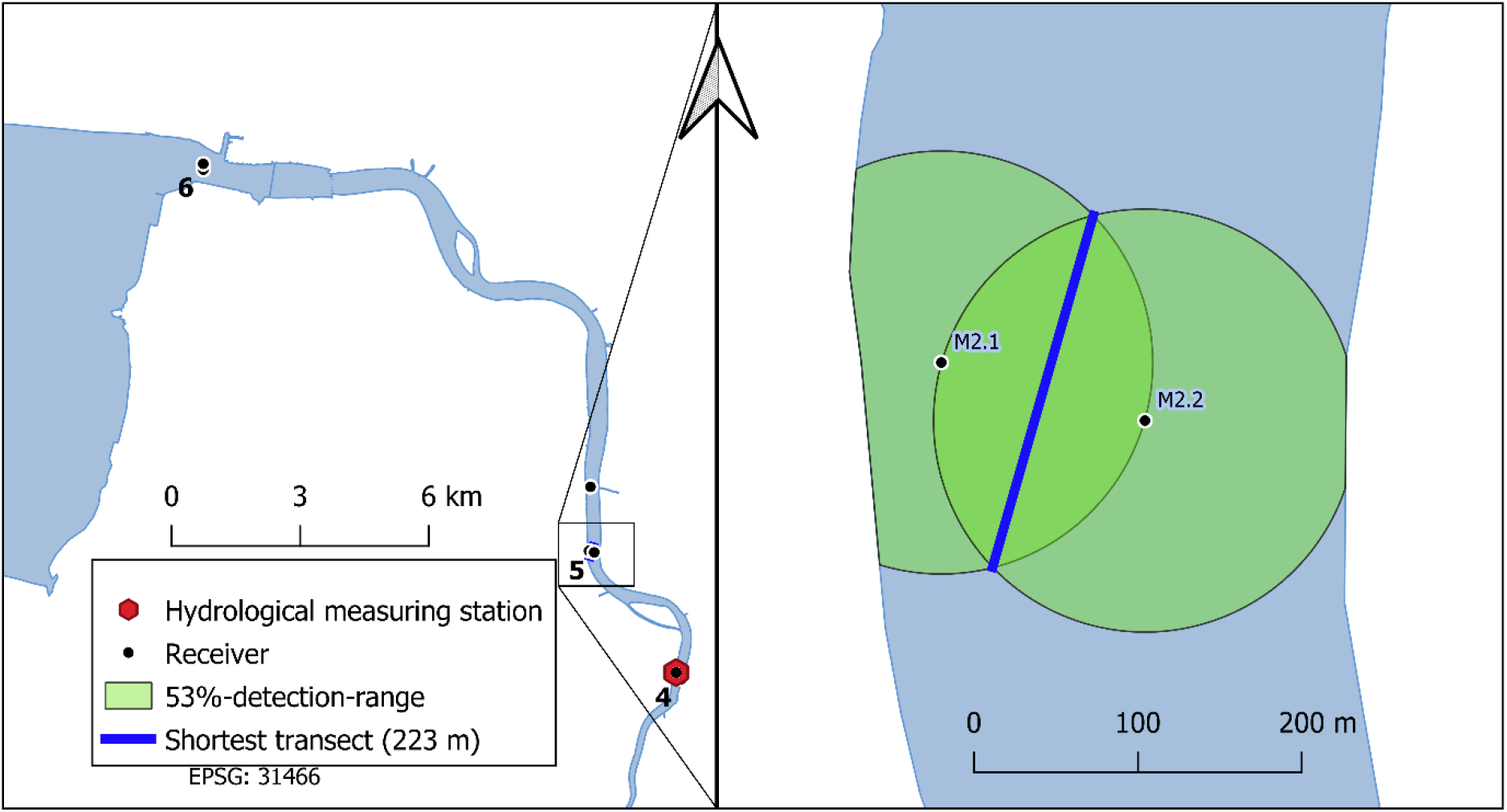
The calculated detection probability of the range-tested setup (Array 5) in the tidal area with the marked 53%- detection-range and the shortest overlapping transect.

**Figure 4:**
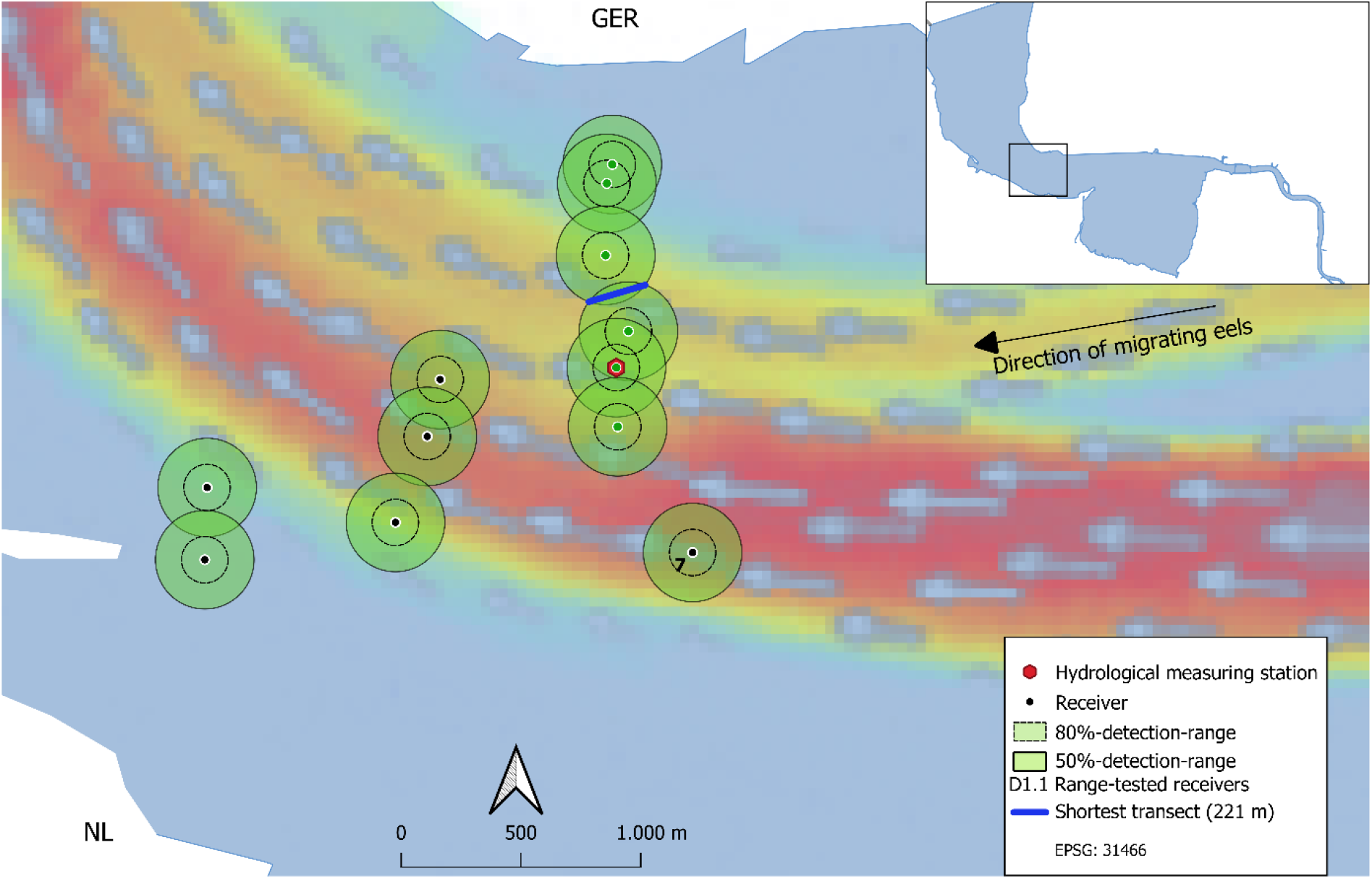
Calculated detection radii during the main eel migration season in autumn of the estuarine receivers (Array 7); 80%- detection range = 98 m; 50%-detection-range = 197 m. The blue arrows indicate the current direction and the colouration symbolize different maximum ebb current velocities (red – fast (∼1.4m/s), yellow – medium (∼1 m/s), green – medium slow (∼0.7 m/s), blue- slow (<0.5 m/s) originating from Herrling et al. (2014).

**Table 4:**
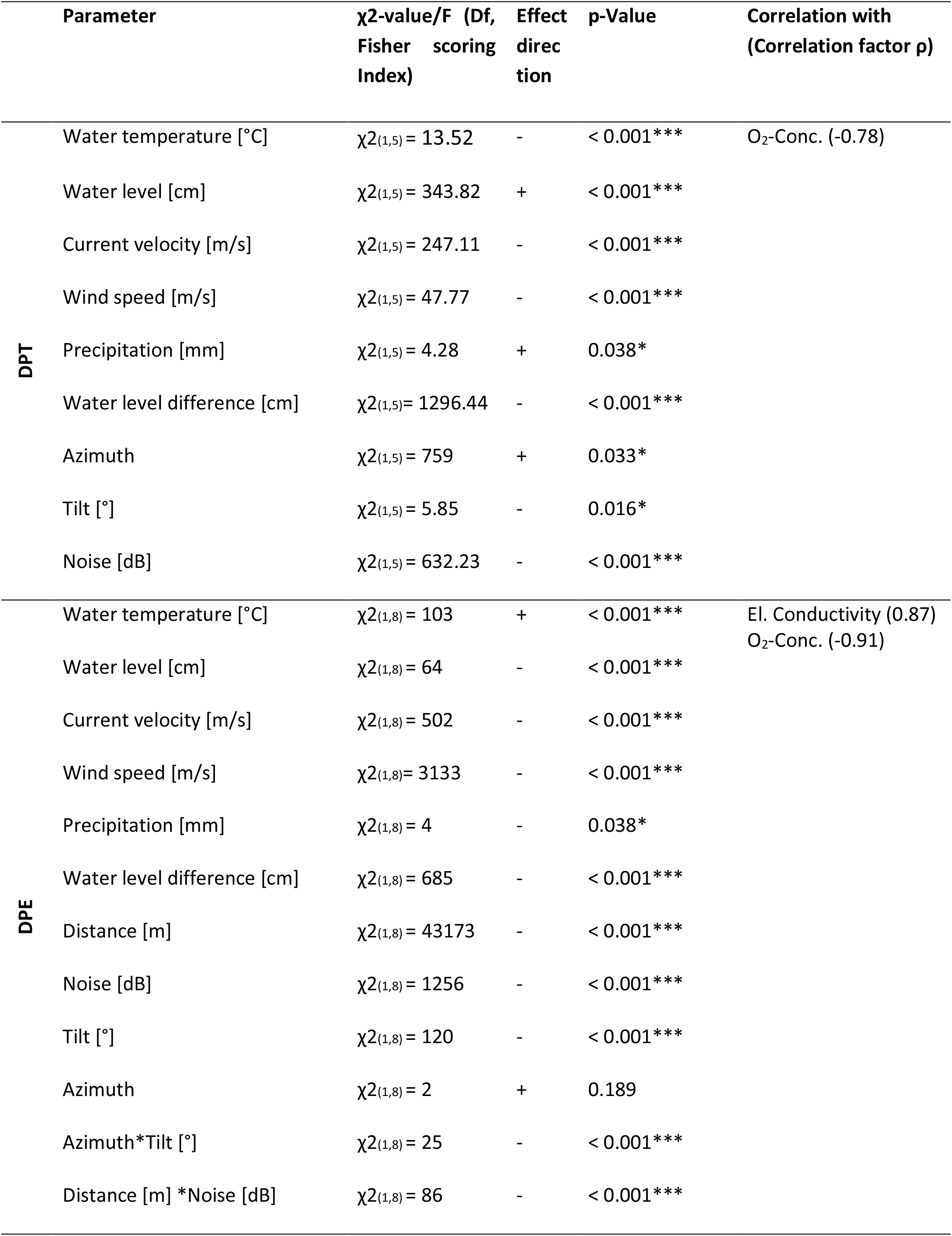
Summary of the minimum adequate models of factors influencing the receiver performance i.e., detection probability in the tidal area and estuary

High current velocities, precipitation, wind speeds, water level and water level differences reduced the detection probability, while warmer water temperatures increased the detection probability in the estuary. Furthermore, a significant negative effect of the tilt angle in dependence of the calculated azimuth was evident. The negative effect of noise on detection probability became stronger with increasing distance (Pseudo-McFadden-R²= 0.591, model “DPE”, Table 4).

### Chance of Detection

In the estuary the majority of eels (92.49%) was detected at receivers of Array 7, covering the main current (based on models by Herrling et al., 2014) of the Ems River (receivers D1.1-D1.6, Figure 1). Here, the shortest transect through the overlap of two receivers’ 50%- detection-range measured 221 m (Figure 4). At average swimming speed, 5-6 pings could be emitted, resulting in a chance of detection of 98.3% under average conditions. In the tidal area the shortest transect within the main current was 223 m (Figure 3) resulting in a chance of detection of 96.3% at average eel swimming speeds (Table 5). These were minimum estimates as the power output of the used V9-acoustic tags was higher (146dB) than the output of the receiver’s sync tags used to determine the reported detection ranges (142 dB). The estimated chances of detection suggested a very high detection efficiency of the setup.

**Table 5:**
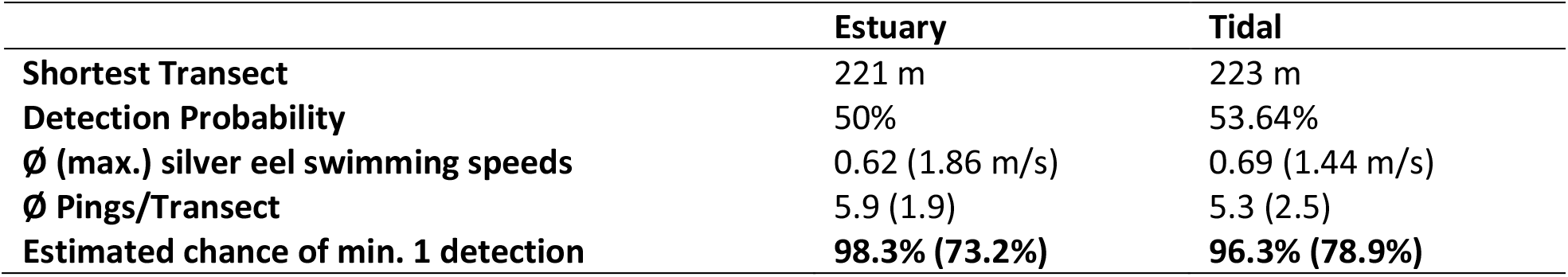
Chance of detection in the tidal area and estuary of observed receiver arrays.

### Detectability

The dominance analysis of the three models per river area revealed that current velocity was a major influencing factor in all models. 97.5% and 78.5% of the eel’s movement speed in the tidal and estuary area, respectively, can be explained by current velocity (which was highly correlated with differences in water level). Further, it accounted for 33.8% and 18.2% of the variation in migration probability in the tidal and estuary area, respectively. Simultaneously, current velocity was a major influencing factor on detection probability in both river sections (relative importance in DPT model: 15.1%; DPE model: 10,0%), as was water level difference in the tidal area (relative importance in DPT model: 19.0%). A minor positive effect of precipitation on detection probability the (relative importance in DPT model: 0.8%), migration speed (relative importance in MST model: 1.1%) and migration activity (relative importance in MST model: 8.75%, here accumulated precipitation) was identified in the tidal area. In the estuary the precipitation had a minor limiting effect on the detection probability (0.2%). Water temperature, however, had a contradictory effect on the detection probability. Higher water temperatures increased the detection probability (relative importance in DPE model: 2.9%) and migration speed (relative importance in MPE model: 4.1%) in the estuary, while decreasing the detection probalility in the tidal area (relative importance in DPT model: 9.5%) (Figure 5 Figure 6).

**Figure 5:**
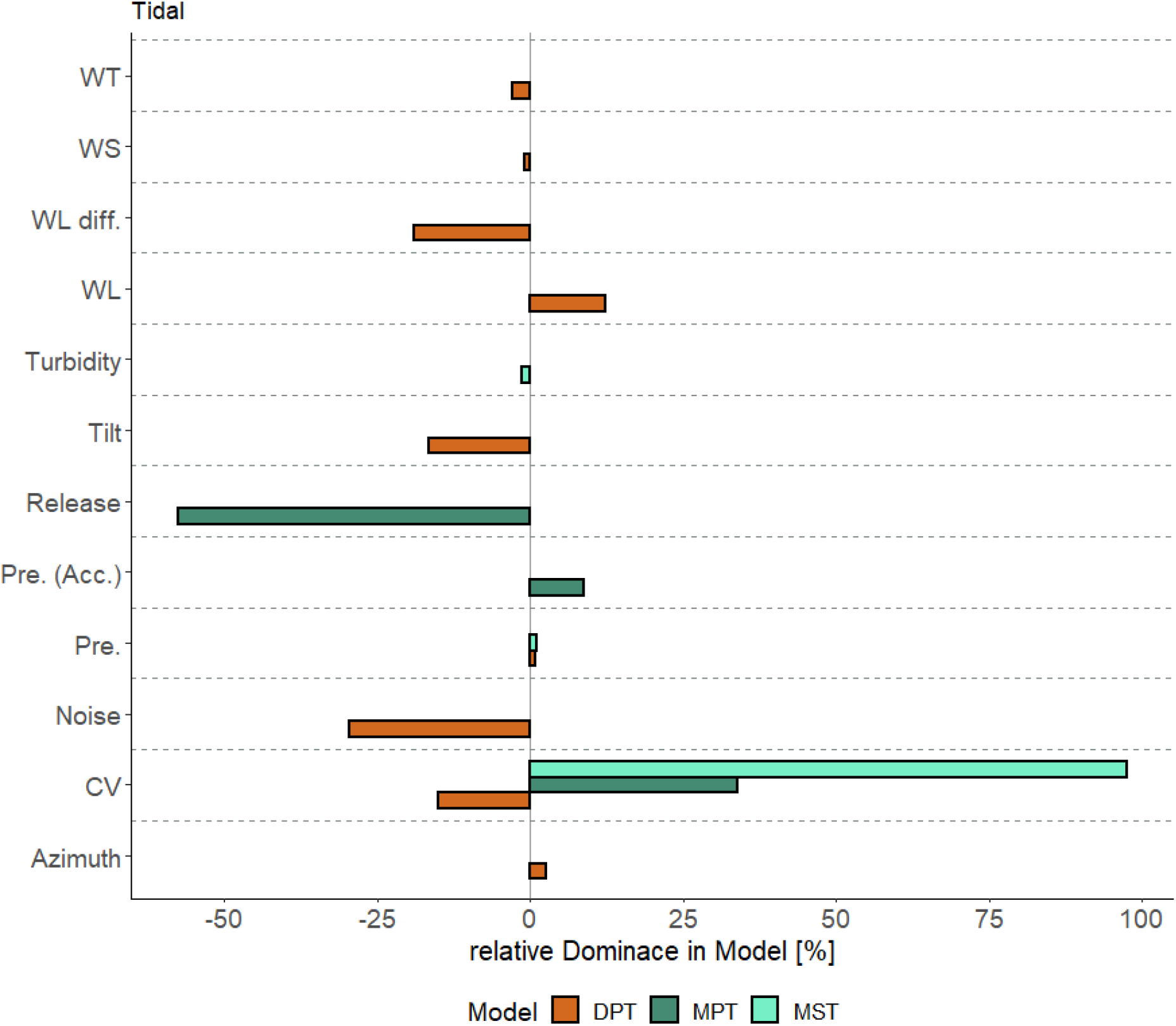
Relative Dominance of the parameters in the final model in the three tested models DPT (Detection probability), MPT (Migration probability) and MST (Migration speed) including their effect direction in the tidal area, with bars to the left indicating a negative effect on the dependent variable and bars to the right indicating a positive effect on the dependent variable. WT- Water Temperature, WS- Wind Speed, WL diff.- Water level difference, WL- Water level, Pre.(Acc.)-Precipitation (accumulated), CV- Current velocity

**Figure 6:**
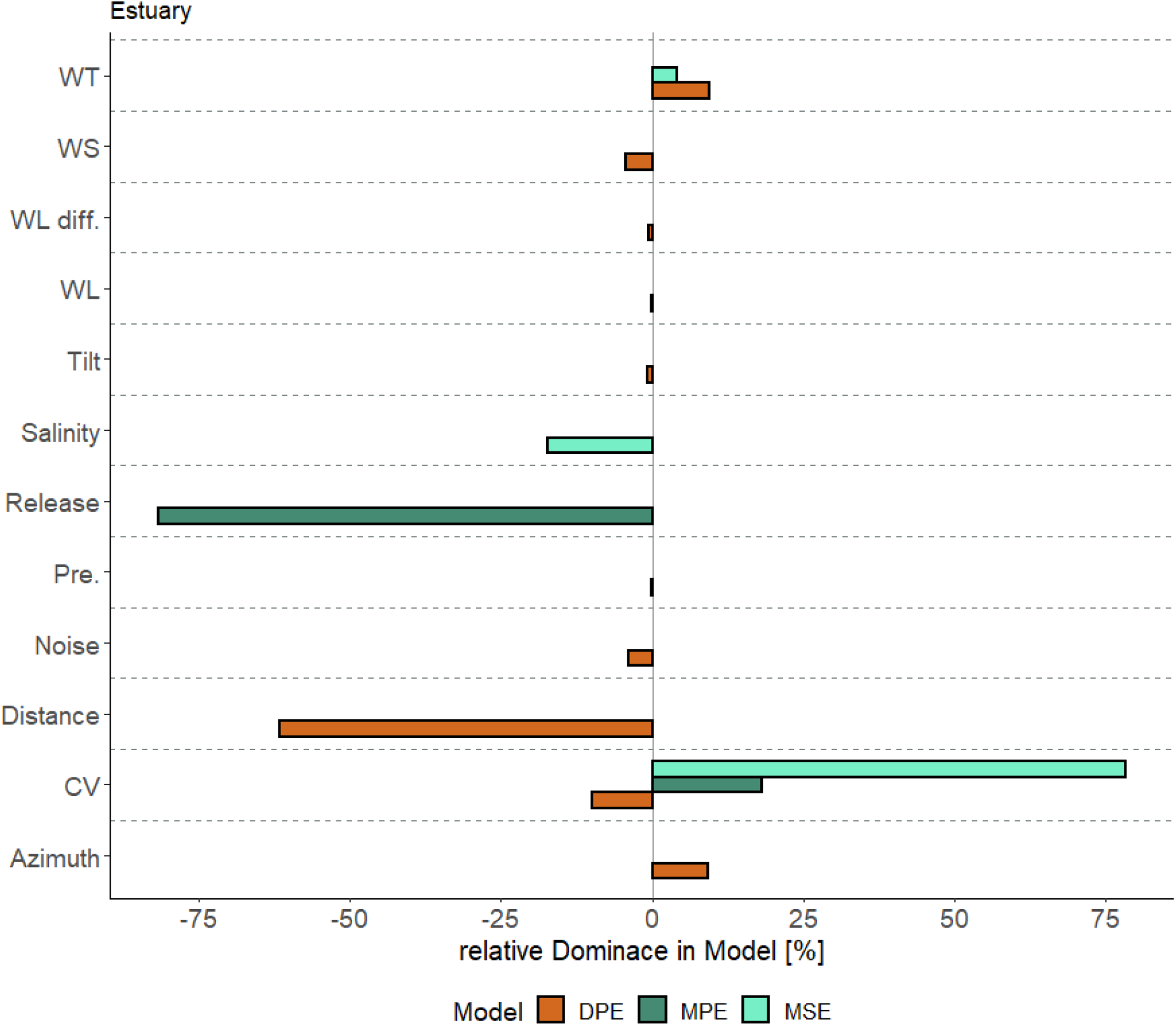
Relative Dominance of the parameters in the final model in the three tested models DPE (Detection probability), MPE (Migration probability) and MSE (Migration speed) including their effect direction in the estuary, with bars to the left indicating a negative effect on the dependent variable and bars to the right indicating a positive effect on the dependent variable. WT- Water Temperature, WS- Wind Speed, WL diff.- Water level difference, WL- Water level, Pre.(Acc.)-Precipitation (accumulated), CV- Current velocity.

## Discussion

Environmental factors are known to influence the activity and progression speed of migrating fish. Acoustic telemetry is a common tool to the investigation of fish migration ecology, but its performance and thus the resulting data quality are not less reliant on the environment. This study assessed the relative influence of environmental factors on both the migration activity and speed of European silver eels, as well as acoustic detection ranges, thereby identifying bottleneck mechanisms to their detectability within acoustic tracking studies.

### Migration Speed and Activity

Average migration speeds in this study were measured at 0.69 ± 0.41 m/s and 0.62 ± 0.44 m/s in the tidal and estuary, respectively. This coincides with observations in the Meuse River averaging at 0.63 m/s (Verbiest et al., 2012) and is slightly higher than silver eels in the non-tidal area of the Rhine River, measured at 0.5 m/s (Breukelaar et al., 2009), while slightly lower than migrations speeds in the tidal area of the Westerschelde with 0.95 m/s (Verhelst et al., 2018). The maximum observed migration speeds of 1.44 and 1.86 m/s in the tidal and estuary areas, respectively, are slightly lower than those recorded by Breukelaar et al. (2009) with 2.20 m/s and coincides with observations in the tidal area by Verhelst et al. (2018) with 1.87 m/s.

Current velocity was the major environmental factor influencing migration activity and speed of silver eels in the tidal and estuary section of the Ems River. Current velocity was positively related to both. activity and speed, which is consistent with other studies (Vøllestad et al., 1986; Cullen & McCarthy, 2003; Bultel et al., 2014; Marohn et al., 2014; Reckordt et al., 2014; Stein et al., 2016; Trancart et al., 2018). In both river sections the eeĺs swimming speed mirrored the current velocity closely, especially during periods of fast flow. This implies a rather passive locomotion, drifting with tidal currents, as also observed by Lenihan et al. (2020). Higher relative migration speeds (i.e., swimming speed corrected for current velocity) only occurred during stagnant waters xof high and low tide peaks. During these tidal phases, active locomotion is the only option to progress migration, even though under higher energy expenditure. The migration activity was highest during downstream currents, with 89.4 and 87.9% of migration events occurring during ebb tide in the tidal and estuarine areas, respectively. In the estuary 92.5% of eels were first detected (per array) by receivers covering the main current. These findings strongly imply the utilization of selective tidal stream transport (STST) by silver eels during their riverine downstream migration. This is in accordance with findings of Verhelst et al. (2018) in the Schelde river. This behaviour likely is an adaption to conserve energy during the initial phase of the long and energy sapping reproductive journey of eels and is a safe strategy to reach the ocean, minimizing risky of wrong turns or detours.

The results showed that higher water temperatures favoured higher migrations speeds in the estuary. All migrations occurred at temperatures between 4.6°C and 16.2°C, while the upper temperature limit above which eel activity diminishes due to low oxygen availability or other metabolic mechanisms was not present during our observation period (Claësson et al., 2016). These temperature related migration preferences fall in range of past observations with 10° - 16°C in Spain (Lobón-Cerviá & Carrascal, 1992), 4° - 18°C in the River Imsa (Norway) (Vøllestad et al., 1986), 8° – 16°C in the Elbe (Germany) (Stein et al., 2016) and 6° - 15°C in the River Loir (France) (Bruijs & Durif, 2009). Corresponding with past studies, the results if this investigation indicate that a continuous drop in water temperature may be influential on the onset and end of the main migration season, after which eels enter winter dormancy, as the ectothermic metabolism limits the capability of eels to be active during cold conditions (Westerberg & Sjöberg, 2015). Movement of eels can then restart again in spring, when temperatures rise above the threshold (Aarestrup et al., 2008; Trancart et al., 2018; Økland et al., 2019). With respect to rising temperatures in the face of climate change this could lead to a continuation or shift of the migration period towards the winter months (Arevalo et al., 2021). Higher ambient temperatures in early autumn may extend the inactive period, while milder winters provide suitable migration conditions. Migration speed in the estuary declined with increasing salinity. This could be linked to tidal changes, as an influx of sea water into the Dollart estuary is represented by elevated salinity levels, opposing the movement direction of eels, which results in lowered migration speeds. During the downstream movements of silver eels, the salinity gradient in tidal and estuarine river areas may support eels in their orientation (Béguer-Pon et al., 2016).

Additionally, eels in the tidal area showed increased activity during prolonged periods of higher precipitation and exhibited higher migration speeds. Periods of high precipitation possibly indirectly enabled faster migration speeds by elevating water level and current velocity in rivers providing optimal conditions for the downstream movement of silver eels. A positive effect of precipitation on the downstream migration of silver eels is supported by studies of Durif et al. (2003), Stein et al. (2016) and Trancart et al. (2018).

Increased turbidity is usually linked with increased activity, as a probable predator avoidance adaptation. Further, increased turbidity hinders the influence of other extrinsic light, such as solar and lunar illumination and allows for regular migrations during day time (Bruijs and Durif, 2009, Monteiro et al., 2021). In this study; high turbidity levels were associated with lower migration speeds in the tidal area. This might be linked to a dilution effect during periods of high river discharge, which creates more favourable migration conditions for many fish (Verbiest et al., 2012). Additionally, the highly anthropogenised river section in the tidal area is prone to extreme turbidity (>1000 NTU), regularly promoting hypoxic or even anoxic conditions (Schulz et al., 2021; Talke & de Swart, 2011) and thus creating barriers for a continuous and fast eel migration (Bultel et al., 2014). These environmental conditions are already reflected by eel behaviour, a species known to be tolerant for low oxygen concentrations (Dean & Richardson, 1999). However, infection with the parasite *Anguillicola crassus* increases the sensitivity for hypoxic conditions. As past studies showed a high infection rate of eels in the area (Lefebvre et al., 2007), hypoxic areas can slow down and possibly decrease the escapement sucess of the downstream migration of silver eels in the tidal area of highly anthropogenised rivers. Subsequently, the migration behaviour of other fish species, such as trout and salmon in the Ems River is expected to be impaired severely by these environmental conditions (Dean & Richardson, 1999).

Apart from the environmental effects discussed above, the migration activity was strongly influenced by the time of release with most eels continuing their migration shortly after release. A similar effect was observed in past telemetry studies (Trancart et al., 2018), highlighting the importance to account for this effect in models to avoid a release bias and therefore a misinterpretation of other parameters. All in all, the models presented in this study predicted the migration speed, migration activity and detection probability well over the observed time period, explaining between 24 and 69 % of the observed variation. As an exception, migration activity in the estuary was poorly explained by the considered factors (Pseudo-McFadden-R²=0.022, model “MPE”). This might be in part due to a lower number of eels (n=173) creating a limited number of migrations events over 1080 tidal cycles. Further, the effect of turbidity could not be tested in the estuary, due to missing and erroneous measurements. The migration activity of downstream migrating eels was probably influenced majorly by factors in the upstream river areas, while in the estuary only tidal influences played a role in the continuation of the reproductive migration.

### Receiver performance

Over the observation period the detection probability in the tidal area at a distance of 127 m was at 53%, while the range at which 50% of all pings were received in the estuarine setup was at 197 m. However, these are slightly lower detection ranges compared to the ground-moored coastal Belgian telemetry network (∼232 m), as proposed by Reubens et al. (2019), due to a shallower receiver installation at buoy chains. Therefore, surface related disturbances (wind and waves), precipitation and the ambient noise of buoy chain rattling can have a comparably larger effect on the receiver performance than in a bottom installed receiver network (Reubens et al., 2019). The latter comes with a cost of complicated accessibility and sedimentation. Therefore, a receiver network installed closer to the surface allows for more regular maintenance, like battery exchange and clearing from biofouling, avoiding a further decrease in receiver performance.

Receiver performance, measured as the detection probability at a given distance, changed under varying environmental conditions. In the tidal river section, water level, precipitation and azimuth increased the acoustic detection range, while high current velocities, ambient noise, tilt, water level differences, wind speeds and water temperatures decreased the chance of receiving pings. In the estuary the receiver performance was limited by high current velocities, precipitation, wind speeds, water level, water level differences and low water temperatures. Additionally, the negative effect of noise increased with rising distance between the receivers, while also a significant negative effect of the tilt angle in dependence of the calculated azimuth was evident.

Current velocity and water level difference were the major environmental impacting factors, negatively influencing the receiver performance in both the tidal and estuarine section. These findings are in accordance with other telemetry range test studies (Melnychuk, 2012; Mathies et al., 2014; Vemco, 2016; Reubens et al., 2019; Rechisky et al., 2020), suggesting that an increased water movement causes higher background noise and hinders signal detection.

Signals from tags and other receivers are more likely to be masked or not recognised under loud conditions. Also, anthropogenic sound sources, such as ship traffic or route maintenance work, were expected in the area, as the receiver ranges in many cases covered nautical navigation routes. In addition, the rattling of buoy chain elements was a major noise source close to the receivers.

Wind speed had a negative effect on detection range in both river sections. Wind induces wave movement (Reubens et al., 2019), trapping air bubbles in the upper water column, thus deflecting and scattering acoustic signals (Gjelland & Hedger, 2013; Kessel et al., 2014). Further, as all receivers were located close to the water surface, the noise of breaking waves may also have been influential. The dominance analysis revealed that the explained variance of detection probability by wind decreased with distance from the river estuary. This coincides with a more sheltered position of the tidal setup and the common meteorological patterns as wind speed was also more extreme in coastal regions than in inland regions. Generally, wind is considered to be an important influencing factor hindering signal transmission in many telemetry studies (Gjelland & Hedger, 2013; Stocks et al., 2014; Kessel et al., 2014; Reubens et al., 2019).

Water temperature had an opposing effect on the detection probability in the tidal and estuarine setup. Normally, higher water temperatures enhance sound transmission and therefore detection probability (Yuwono et al., 2014). However, the presence of algae is also indirectly linked to higher water temperatures in early autumn, which can interfere with the sound propagation in water (Winter et al., 2021). The observation period was set to cover the main migration period of the year which coincides with the roughest environmental conditions in the area. Therefore, the environmental conditions for the observation of summer migrating fish with acoustic telemetry are probably less limiting.

### Dominance Analysis and Detectability

The dominance analysis approach allowed to rank order and compare the relative influence of environmental factors on the migration behaviour of eels and the acoustic detection range. Current velocity, closely correlated with differences in water level, had a major positive influence on migration activity and speed in both the tidal and estuarine area, explaining between 18 and 98% of the observed variance in these contexts. On the other hand, current velocity and water level difference were a major influencing factor limiting the receiver performance in both tested areas, explaining between 10 and 19% of the observed variances. Therefore, this factor creates a bottleneck for detectability during the main migration phases. This is a highly relevant finding as it is likely transferable to other migrating fish species in tidal and estuarine rivers that use STST. Therefore, when studying these species emphasis should be put on careful receiver placement in areas with high current velocities to avoid underestimating migrations during conditions of high or regular tidal currents. Single-time surveys of detection range on a newly installed telemetry setup, e.g., through boat drift tests, should be carried out during periods of high river discharge to obtain the minimum effective detection range.

Similarly, but to a much smaller extent, precipitation increased migration activity and speed in the tidal area. At the same time, precipitation limited receiver performance in the estuary (albeit with a relative minor importance). Further, higher water temperatures enabled higher migration speeds in the estuary while simultaneously enhancing the sound propagation (Yuwono et al., 2014; Makar 2022) favouring successful signal transmission. Conditions of high salinity in the estuary, closely correlated with electric conductivity water temperature, create beneficial conditions for detecting migrating eels because eels are slowed down and signal transmission is improved as the higher density of water benefits sound propagation (Yuwono et al., 2014; Makar 2022). Notably turbidity can also be a limiting factor regarding detectability, which was not the case in this study. However, moderate turbidity levels can favour eel activity (Bruijs and Durif, 2009; Monteiro et al., 2021), while a higher number of particles in the water column can scatter acoustic signals (Singh et al., 2009; Vemco, 2016).

## Conclusion

This study suggests that European silver eels use selective tidal stream transport (STST) and passive locomotion during their downstream migration through tidal rivers. Likewise, current velocity was identified as the major driver of migration activity and speed in tidal and estuarine river landscapes. However, current velocity was also a major factor limiting the detection range of acoustic telemetry systems. As many migratory fish species use STST, this bottleneck can lead to underestimations of movement activity in telemetry studies during high currents, often coinciding with main migration phases. Therefore, this study emphasizes a careful placement of acoustic receivers in areas with regular strong tidal currents, especially when studying fish using STST.

## Supporting information

Supplemetary file (with tables, texts and graphs)

## Acknowledgements

We want to acknowledge Ulrike Wypler for her invaluable help with conducting the field work. Appreciation also goes to Martin Goldsweer for operating the catch gear . Permissions to install the acoustic receiver network were generously provided by the local water authority (WSV Ems-Nordsee). We greatly appreciate the support of Thomas Schlüsselburg (WSV Ems-Nordsee) and crews from the buoy tenders “Norden”, “Friesland” and “Gustav Meyer” with receiver mounting and maintenance. This study was conducted as a part of the project “BALANCE”, for which financial support was provided by the Lower Saxony Ministry of Food, Agriculture and Consumer Protection and the European Maritime Fisheries Fund (EMFF-ID: NI-1-18-004). Some of the acoustic receivers deployed in the Dollart estuary were part of the “Ruim Baan voor Vissen” project, financed by Waddenfonds (01755849 / WF- 2019/200914).

