## Supplementary material for "To hear or not to hear: How Selective Tidal Stream Transport Interferes with the Detectability of Migrating Silver Eels in a Tidal River": Supplemetary file (with tables, texts and graphs)

### Appendix

*Table A1: Distance matrix of the estuarine setup*

| Distance (m) | D1.1 | D1.2 | D1.3 | D1.4 | D1.5 | D1.6 |
|---|---|---|---|---|---|---|
| D1.1 | - | 85 | 385 | 707 | 855 | 1097 |
| D1.2 | | - | 300 | 631 | 779 | 1022 |
| D1.3 | | | - | 331 | 479 | 722 |
| D1.4 | | | | - | 172 | 409 |
| D1.5 | | | | | - | 242 |
| D1.6 | | | | | | - |

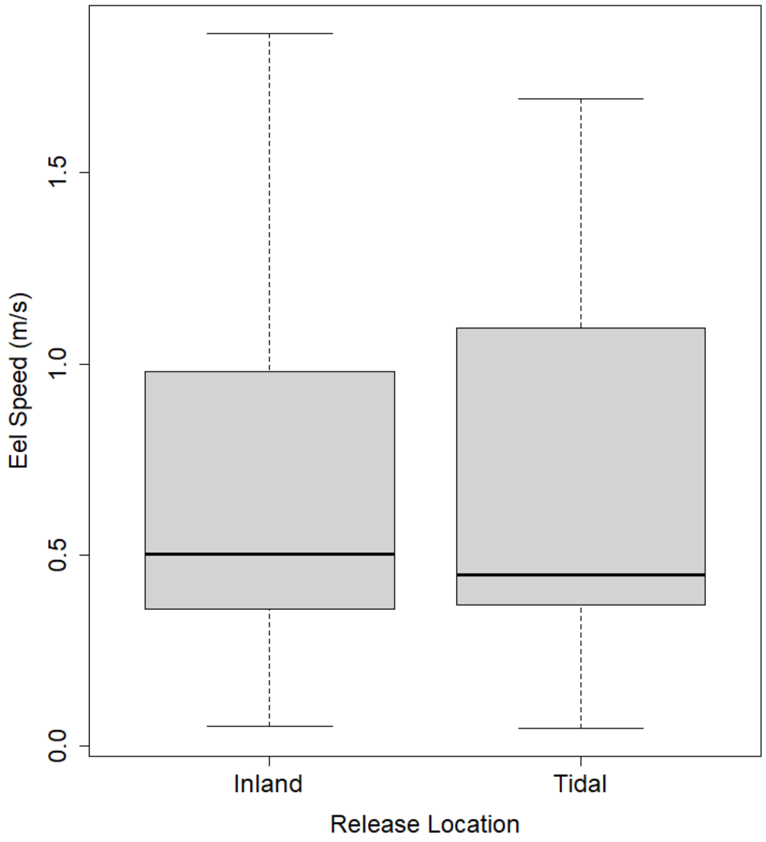


*Figure A1: Boxplot of the Swimming speeds of eels in the tidal area dependent on their release location.*

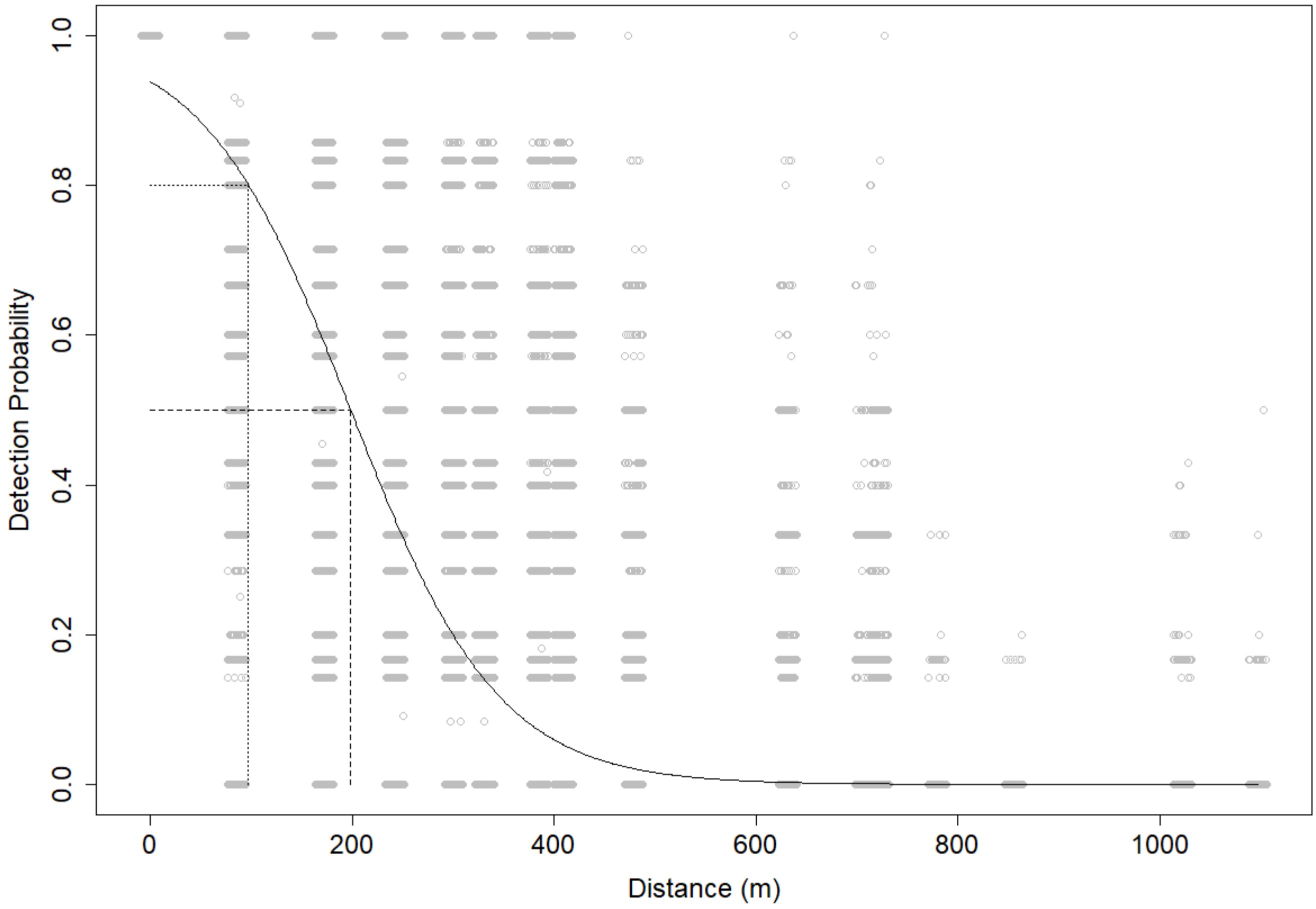


*Figure A2: The influence of distance in the detection probability in the estuary with marked 50% and 80% detection range (197 m and 98 m respectively)*

*Table A2: Summary of the R² values of the model and the respective variables of the final model for the movement speed and migration activity*

| Parameter | McFadden-Pseudo-$R^2$ *Nagelkerke- Pseudo-$R^2$ | Relative Importance [%] |
|---|---|---|
| **MST** | 0.545* | 100 |
| Current velocity [m/s] | 0.531* | 97.5 |
| Turbidity [NTU] | 0.008* | 1.4 |
| Precipitation [mm] | 0.006* | 1.1 |
| **MSE** | 0.689* | 100 |
| Current velocity [m/s] | 0.540* | 78.5 |
| Salinity [‰] | 0.120* | 17.4 |
| Water temperature [°C] | 0.008* | 4.1 |
| **MPT** | 0.240 | 100 |
| Accumulated Precipitation [mm] | 0.021 | 8.75 |
| Current velocity [m/s] | 0.081 | 33.75 |
| Time since Release [h] | 0.138 | 57.5 |
| **MPE** | 0.022 | 100 |
| Current velocity [m/s] | 0.004 | 18.2 |
| Time since Release [h] | 0.018 | 81.8 |

*Table A3: Summary of the $R^2$ values of the model and the respective variables of the final model for the detection probability*

| Parameter | McFadden-Pseudo-$R^2$ | Relative Importance [%] |
| --- | --- | --- |
| **DPT** | 0.489 | 100 |
| Water temperature [°C] | 0.014 | 2.86 |
| Water level [cm] | 0.06 | 12.27 |
| Current velocity [m/s] | 0.074 | 15.13 |
| Wind speed [m/s] | 0.005 | 1.029 |
| Precipitation [mm] | 0.004 | 0.82 |
| Water level difference [cm] | 0.093 | 19.02 |
| Azimuth | 0.013 | 2.661 |
| Tilt [°] | 0.081 | 16.56 |
| Noise [dB] | 0.145 | 29.65 |
| **DPE** | 0.591 | 100 |
| Water temperature [°C] | 0.056 | 9.46 |
| Water level [cm] | <0.001 | <0.1 |
| Current velocity [m/s] | 0.059 | 9.97 |
| Wind speed [m/s] | 0.026 | 4.39 |
| Precipitation [mm] | 0.001 | 0.17 |
| Water level difference [cm] | 0.003 | 0.51 |
| Distance [m] | 0.365 | 61.66 |
| Noise [dB] | 0.023 | 3.89 |
| Tilt [°] | 0.004 | 0.68 |
| Azimuth | 0.055 | 9.29 |
| Azimuth*Tilt [°] | -- | -- |
| Distance [m] *Noise [dB] | -- | -- |

### RANGE-TESTING OF THE EMS-TELEMETRY-NETWORK

Supervised by:

Prof. Dr. Reinhold Hanel — Thünen Institute of Fisheries Ecology
Prof. Dr. Marko Rohlfs — University of Bremen

Benedikt Merk

Matriculation number: 4244349

### Table of Content


Appendix ... 0

1. Introduction ... 6
2. Material and Methods ... 8
    - 2.1. Study Location ... 8
    - 2.2. The Setup ... 9
    - 2.3. Data Analysis ... 10
3. Results ... 13
    - 3.1. Receiver Test ... 13
    - 3.2. Tidal and Inland Receiver Comparison ... 19
4. Discussion ... 20
    - 4.1. Receiver Range-Test ... 20
    - 4.2. Inland and Tidal Receiver Comparison ... 22
    - 4.3. Study Limitations ... 23
    - 4.4. Outlook ... 24
    - 4.5. Conclusion ... 25
5. References ... 26
6. Additional Material ... 28

### 1. Introduction

Acoustic telemetry is an essential method to track animal movement and behavior over an extended time period. It is based on the transmission of acoustic signals, referred as pings, from an acoustic transmitter tag located in or on an organism’s body to a receiver or hydrophone. Further, telemetry can be subdivided in an active and passive approach. While active telemetry involves the active search of a tagged animal and is therefore limited by time and labor. Passive telemetry is based on designated tracking stations along the animal routes and therefore requires more preparation. However, the latter is more beneficial when observing a higher quantity of organism over a longer time period. For both approaches, knowledge regarding the target organism´ behavior is essential in order to place acoustic receivers at crucial landmarks where the organism can be expected (Kessel et al. 2014; Béguer-Pon et al. 2018b).

To allow answering research questions in more detail, especially regarding the animals’ behavior, specific requirements on the telemetry setup need to be met. Therefore, understanding the telemetry system in its environment is crucial to allow for an accurate data interpretation (Kessel et al. 2014; Crossin et al. 2017; Béguer-Pon et al. 2018b).

The detection range of transmitters by each receiver in a common system is dynamic and may depend on a number of different environmental factors. Various studies mention currents and turbidity as main influential factors (Mathies et al. 2014; Reubens et al. 2019). Furthermore, biofouling on the receivers, weather events such as storms as well as swell have been reported of having potential negative impact on the detection range of acoustic receivers (Kessel et al. 2014; Mathies et al. 2014; Reubens et al. 2019) . Additionally, ambient noise sources with anthropogenic (nautical traffic), biotic (e.g., crabs, fish) or abiotic (e.g. waves, currents) origin are known to potentially conceal transmitter pings (Mathies et al. 2014; Reubens et al. 2019). Moreover, some studies identified topographical irregularities interfering with acoustic signals by causing shadow or scattering effects and bathymetric influences (Selby et al. 2016; Crossin et al. 2017; Babin et al. 2019). All these environmental influences are expected to differ largely between inland and tidally influenced areas.

In order to support a comparable validation of the telemetry system the minimal detection range and/ or the 50%-detection range are commonly stated in telemetric studies (Kessel et al. 2014; Selby et al. 2016). The latter is defined by the radius where half of all emitted pings are detected. The minimal detection range in this study is referred to a range in which a 95%-detection probability is provided (cf. Whitty et al. 2009).

The here presented telemetry network, was installed within a PHD- project called “BALANCE”, which was initiated to quantify numbers of migrating silver European eels (*Anguilla anguilla*). As the migratory behavior of *A. anguilla* is not completely understood, passive acoustic telemetry can provide valuable insights into the undescribed behavioral patterns (Béguer-Pon et al. 2018b). Earlier studies investigating eel escapement (Breukelaar et al. 2009) calculated average swimming speeds of 0.07 up to 0.93 m/s for different silver eel maturation stages in the Rhine river. Here, the maximum recorded speed was 1.7 m/s. In the river Meuse an average migration speed of 0.62 m/s with a maximum of 1.93 m/s was observed (Verbiest et al. 2012). An average migration speeds of 0.42 m/s were recorded in the river Schelde (Westerschelde), with an average tidal migration speed of 0.95 m/s (Verhelst et al. 2018). All above mentioned swimming speeds solely accounted for females, as implanting transmitters into male eels provide major challenges due to their smaller size. However, their shorter general body length suggests slower migration speeds, as the swimming speed is thought to be linked to the eels' body length (Palstra et al. 2008; Lennox et al. 2018).

For migrating eels in this study an average speed of 0.95 m/s (Verhelst et al. 2018) and a maximum migration speed of 1.93 m/s (Verbiest et al. 2012) is assumed, since eels occasionally use the river discharge as support (Breukelaar et al. 2009; Marohn et al. 2014). This transfers into travel distances of 57 and 116 meters during one 60 second ping interval respectively.

For the BALANCE-Project an acoustic telemetry network was installed in the German river Ems to monitor the behavior and quantification of migrating silver eels (*Anguilla anguilla*). In order to achieve this, acoustic transmitters (Model V9, Vemco Ltd. Halifax, Canada) are carefully implanted into the body cavity of wild-caught migrating silver eels. These tags have a ping-transmission frequency of 60 seconds. To ensure the possibility of detection, it needs to be assured that receiver stations are located reasonably, so that tagged eels can pass the receivers during their downstream migration. Additionally, it needs to be confirmed that at least one ping can be recognized inside the receiver's detection field while the eels are passing by. This implies the minimal detection range of each receiver should exceed the range an eel is able to travel with in the 60 second ping interval.

This study presents results from conducted range tests, that are of great importance to validate the quality of the installed receiver network and to maximize the potential detectability of tagged eels within the BALANCE project. Further, tidal influence on the telemetry network is tested as a summary of a variety environmental factors. This leads to the following research questions and hypothesis:

<u>Research question 1</u>: Can tagged eels be reliably detected by the range-tested receivers?

Hypothesis 1: The 95%-detection range for each receiver is at least 116 m.

Research question 2: Is the receivers detection range impacted by tidal influence?

Hypothesis 2: The detection range of inland receivers is exceeding that of tidal receivers.

### 2. Material and Methods

#### 2.1. Study Location

Receivers in the BALANCE telemetry network are located in the Ems river between Meppen and the Dollart estuary. Their primary purpose is to detect tagged European eels (*Anguilla anguilla*), which are migrating from freshwater to the sea. The telemetry network in the Ems river consists of 29 (Model VR2Tx, Vemco Ltd Halifax, Canada) - receivers located at six stations around river branches as well as catch and release sites of the Balance-project (Figure 1).

In this study the detection range of six receivers, that were considered representative for the system, were investigated by boat drifts. The selected receivers were attached to a variety of objects in the river (Table 1).

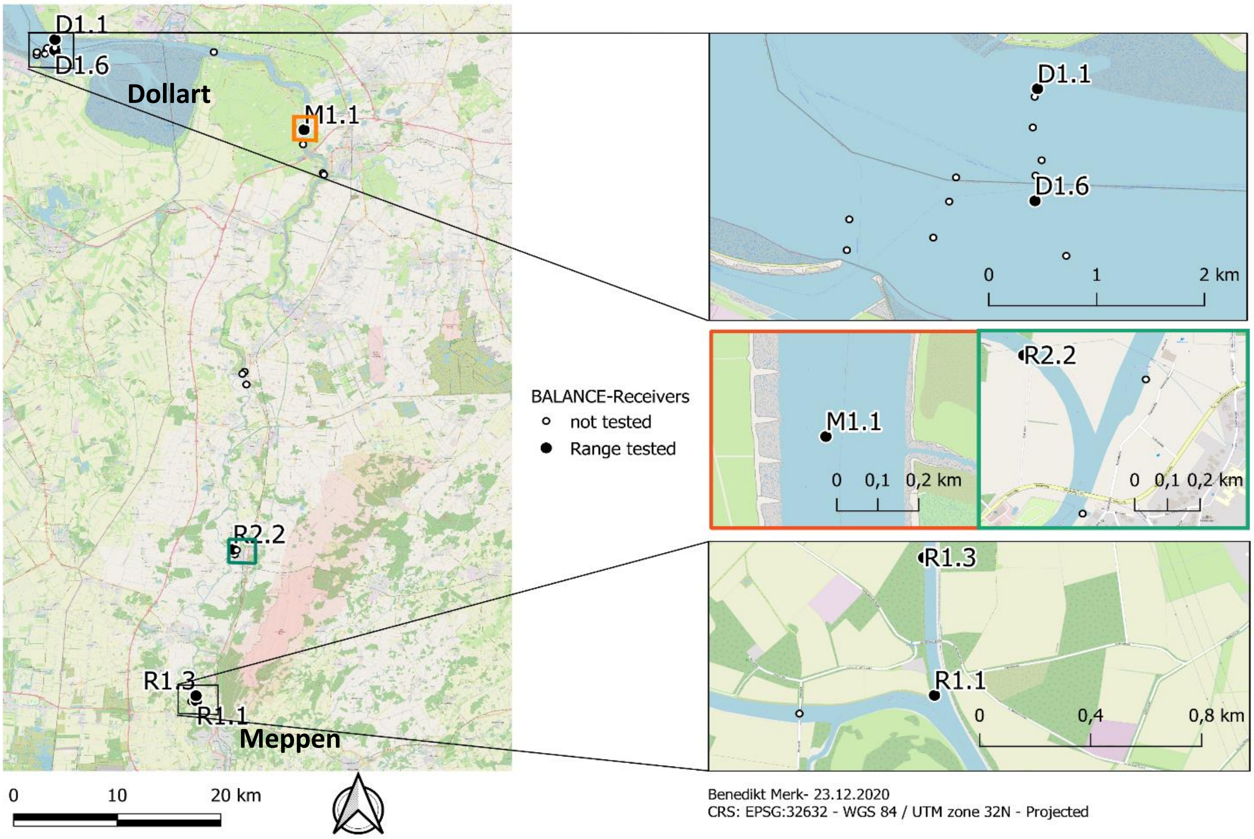


*Figure 1: Receiver locations of the acoustic telemetry network in the Ems-River. North-arrow at the bottom valid for all maps.*

*Table 1: Tested receivers of this study with attachment and river zoning information*

| Receiver | River Zone | Attachment |
|---|---|---|
| R1.1 | Inland | Measuring dolphin |
| R1.3 | Inland | Sheet piling |
| R2.2 | Inland | Fishing berth |
| M1.1 | Tidal | Buoy chain |
| D1.1 | Tidal | Autonomous buoy |
| D1.6 | Tidal | Buoy chain |

Prior to the boat drifts the receivers were adjusted in the range-test setting to achieve a frequent 10 min update of the detected pings (Vemco 2016). A Vemco-V9 Range-test-tag emitted acoustic pings every 23 seconds with 69kHz (Vemco 2020) mimicking the regular V9 tags implanted in the silver eels in regard to power output.

#### 2.2. The Setup

A modified fishing rod with a 250 g weight on a side line was utilized to stabilize the tag in the current at a water depth of 2-2.5 m (Figure 2), to avoid ground contact and minimize losing the tag. A hydrophone (Model VHTx-69kH, undirected, Appendix Figure 10) connected to the VR100-computer was released directly next to the tag in order to detect the emitted pings. Thus, each ping could be assigned to a specific time and location. After each drift the hydrophone was used to communicate with the receiver to examine the detected pings. Additionally, current environmental parameters during the drift, such as tide and swell, were collected as categorial variables. Each receiver was tested in three different drifts. The drifts were conducted at different perpendicular distances of the receiver to maximize the covered area. This allows a more differentiated interpretation, as a near drift can produce other results then a far one due to underwater obstacles. Moreover, the drifts were mostly driven by the current, with some minor exceptions to avoid object collision by small paddling or motor correction. Nevertheless, the third drift at D1.6 was conducted entirely with a running motor on the lowest setting to avert beaching the boat during low tide.

The various environments in which the receivers are stationed required different boat types to enable the drifts. Drifts in the inland zone were conducted with a small rubber boat with a 2PS electric motor. The M1.1 receiver was tested using a hard-shell boat with an integrated motor. Range test for the Dollart receivers D1.1 and D1.6 were conducted employing a rented 200PS rubber and hard-shell combination boat.

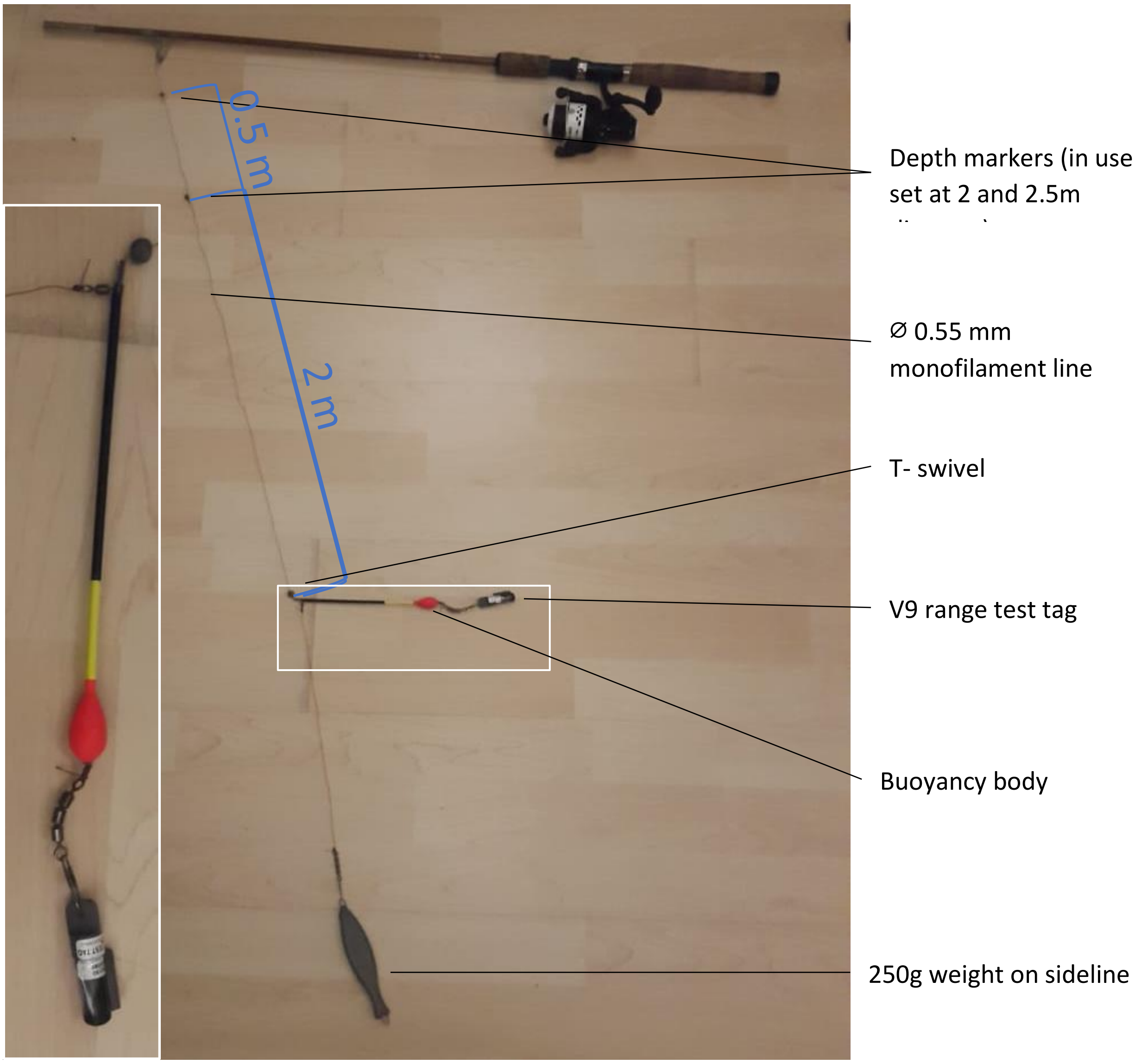


*Figure 2: Setup of the range-test tag on the modified fishing rod*

#### 2.3. Data Analysis

The data of the pings (time and coordinates) detected by the hydrophone were transferred directly from the VR100 (Vemco 2019). The receiver data was accessed by retrieving the receivers after all three drifts were conducted. In case of buoy associated receivers, this needed the support of a buoy tender working vessel, courtesy from the Wasserstraßen- und Schiffahrtsamt (WSA) Ems-Nordsee. Subsequently, the data was manually combined by using the time as reference. Minor delays between VR100 and receiver time were adjusted. Moreover, missing tag pings in the VR100 data were filled up with the according time interval by adding 23 sec to the time of prior ping. In addition, the missing geographic information is calculated by using an arithmetic mean of the prior and later ping coordinates or utilizing a linear function.  These missing pings are caused while the hydrophone was above the water surface due to wave action or checking for entangled drifting plant matter.

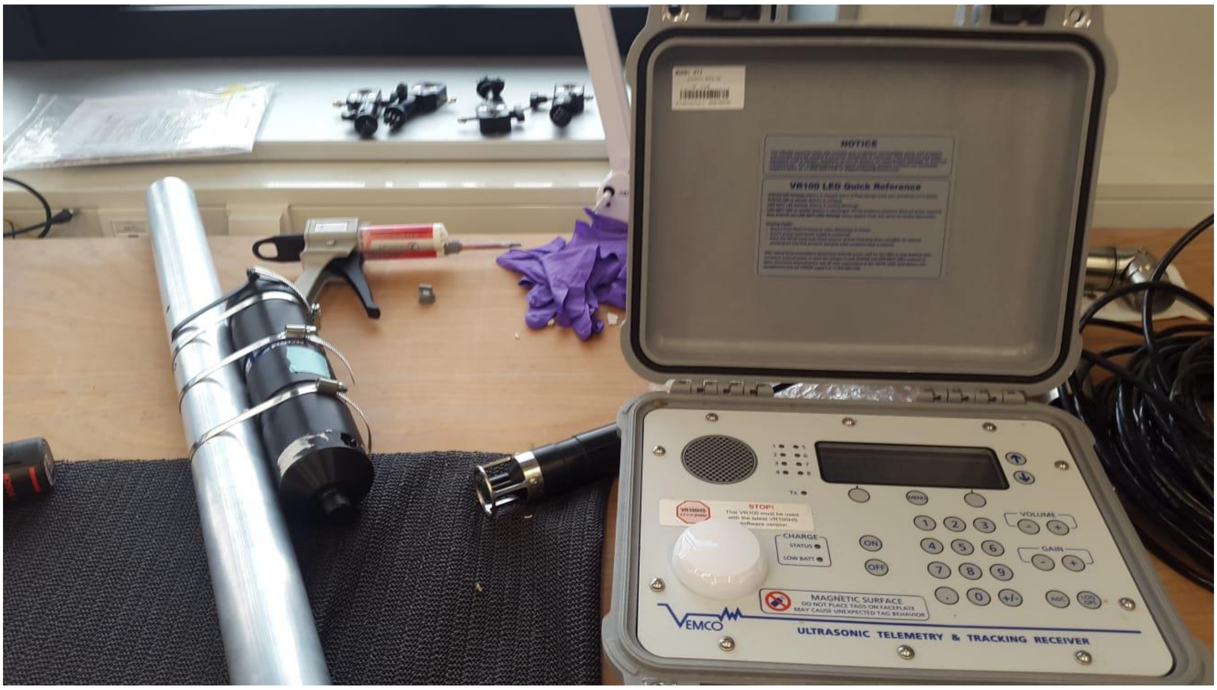


*Figure 3: VR2Tx Receiver attached on a metal pipe (left); VHTx-69kHz hydrophone tip (middle); VR100 (right)*

The coordinates of the pings and of the receiver were transferred to QGIS (Vers. 3.14 “Pi ”) (QGIS.org, 2021) into different layers. The imported data was transformed into a shape file with CRS: EPSG 32632, followed by creating a distance matrix between ping coordinates and receiver location. Eventually, this results in data including the distance of each ping to the receiver with respective binary information about the detection. Subsequently, statistical analysis was conducted with R (Vers 4.03) (R Core Team 2020).  Therefore, a Generalized Linear Model (GLM) with binomial distribution was utilized to test the effect of the distance (general model) as well as drift and distance in combination on the detection probability in separate models (drift model).

The models for the inland receivers were fitted with an additional direction information (upstream, downstream, channel) contingent upon their position at river junctions and/or if they are facing in a particular direction. The attachment object (Measuring dolphin, Sheet piling, Fishing berth; Table 1) is expected to cast acoustic shadows. Therefore, the range in the upstream direction is incomparable to the downstream range. In this case the receiver range diameter is the summation of upstream and downstream radii.

The combined models were analyzed after the Minimal Adequate Models (MAM) approach with a stepwise backward deletion based on a Type-II-Anova to eliminate non-significant variables. The significance threshold for the statistical analysis was set at P= 0.05. The model assumptions were examined visually in regards to the Cook´s distance, Q-Q- and residual plots. A binomial model was accepted if the dispersion parameter was between 0.5 and 2. Otherwise a quasibinomial model was applied.

Consequently, the regression lines of the respective general models were used to calculate the 95%- and 50%-detection-range. These ranges are chosen, since an 100%-detection range is unreliably represented by a binomial model. The absolute maximum-distance is based on the furthest detected ping. Subsequently, buffer areas with the calculated ranges as radii around the receivers were created for visualization with GIS. The created layers were clipped to fit the river shoreline if necessary.

Lastly, the tidal influence on the range was tested by comparing the 50%-detection range of all six receivers. A GLM with gaussian distribution was utilized. Pings opposing the direction of the receiver's hydrophone are excluded, meaning that for inland receivers solely pings in the upstream direction were included.

### 3. Results

The number of emitted pings varied between 28 and 127 over a drifting duration of 12 to 52 min, depending on the current velocity. In total 1031 pings were analyzed of which 649 were detected by the receivers. A significant negative influence of distance on the detectability was validated for all tested receivers (Appendix Table 5).

#### 3.1. Receiver Test

The receiver range differed between the receivers, with R1.3 showing the furthest radius of 95%-detection range in upstream direction with 394 m. In contrast, receiver M1.1 had no modeled 95%-detection range (Figure 4, no 100% certainty at 0 m). All receivers exhibited a 50%-detectability-radius with the largest at 482 m (R1.3 upstream) and the smallest at 96 m (R.2.2 downstream). Generally, the detection range was higher in the direction to which the receiver´s hydrophone was facing. The absolute maximum distance between a detected ping and the receiver was measured at 535 m at receiver M1.1 (Table 2).

Considering the 95%- detection diameter, R1.3 exhibited the largest at 394 m, followed by D1.6 with 347 m. Whereas, the sum of the 95%-detection range for R1.1 added up to 309 m. The smallest inland range was 194 m at R2.2, whilst the smallest total diameter was calculated at 152 m around receiver D1.1 (Table 3).

The calculated ranges are visualized in the following graphic with maps indicating the different probability-radii and the respective model diagrams (Figure 4).

*Table 2: Summary of different detection radii of tested receivers*

| Receiver | Direction | 50%-Range-Radius | 95%-Range-Radius | Max. Range-Distance |
|---|---|---|---|---|
| | | [m] | [m] | [m] |
| D1.1 | NA | 175 | 76 | 242 |
| D1.6 | NA | 260 | 174 | 445 |
| M1.1 | NA | 177 | NA | 535 |
| R2.2 | Upstream | 184 | 129 | 200 |
| | Downstream | 96 | 65 | 103 |
| R1.1 | Upstream | 344 | 265 | NA |
| | Channel | 247 | 43.722 | NA |
| R1.3 | Upstream | 482 | 394 | 485 |
| | Downstream | 175 | NA | 266 |

## D1.1

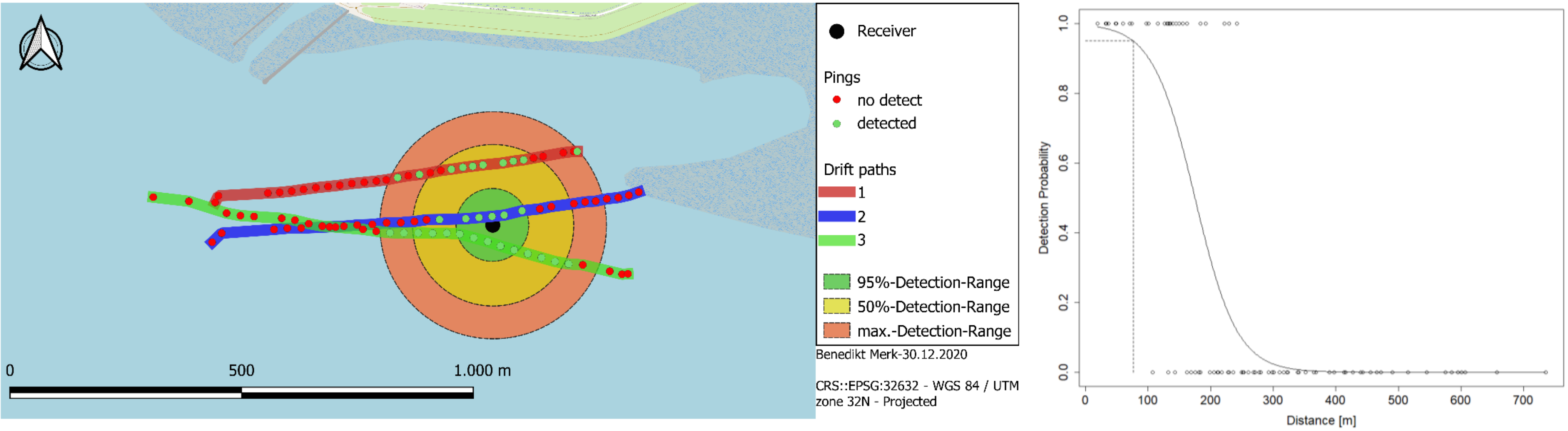
Receiver
Pings
no detect
detected
Drift paths
1
2
3
95%-Detection-Range
50%-Detection-Range
max.-Detection-Range
Benedikt Merk-30.12.2020
CRS::EPSG:32632 - WGS 84 / UTM zone 32N - Projected
0
500
1.000 m
Detection Probability
0.0
0.2
0.4
0.6
0.8
1.0
0
100
200
300
400
500
600
700
Distance [m]

## D1.6

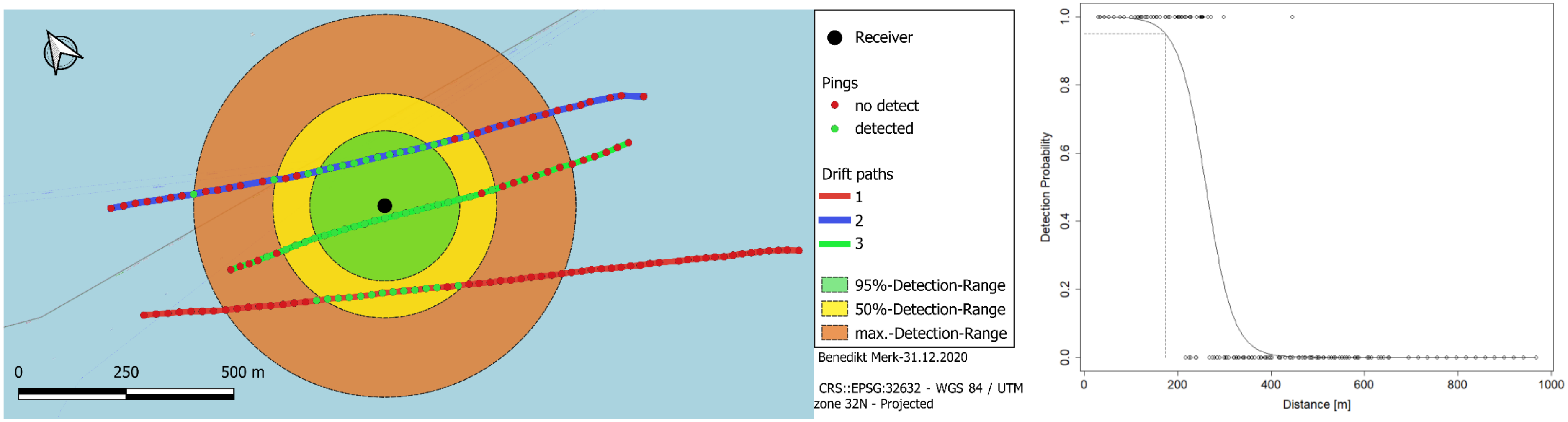
Receiver
Pings
no detect
detected
Drift paths
1
2
3
95%-Detection-Range
50%-Detection-Range
max.-Detection-Range
Benedikt Merk-31.12.2020
CRS::EPSG:32632 - WGS 84 / UTM zone 32N - Projected
0
250
500 m
Detection Probability
0.0
0.2
0.4
0.6
0.8
1.0
0
200
400
600
800
1000
Distance [m]

M1.1

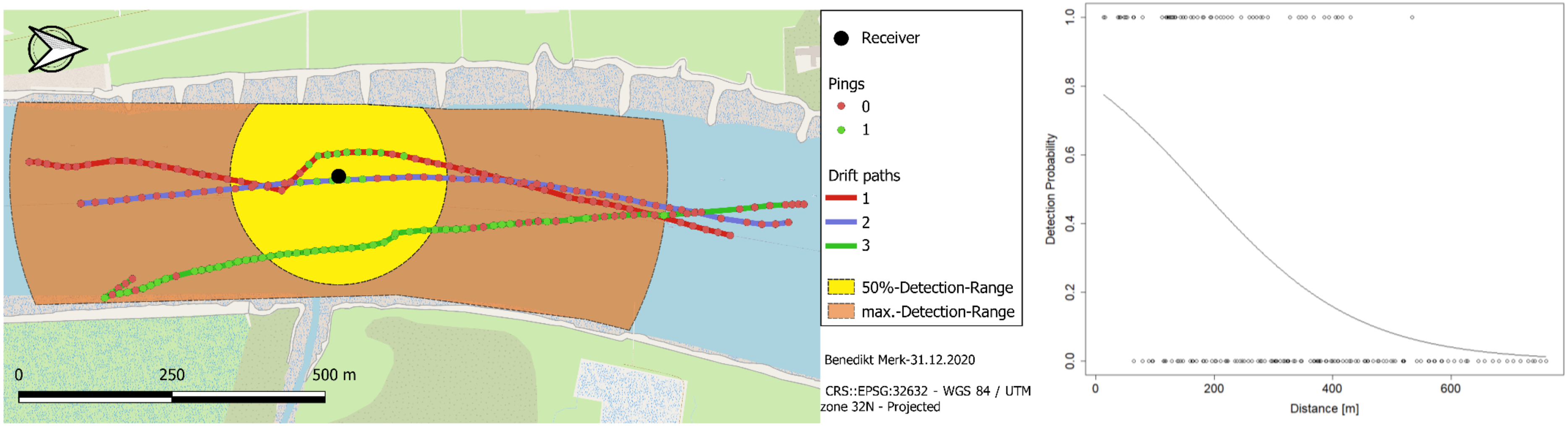

0
250
500 m
Receiver
Pings
0
1
Drift paths
1
2
3
50%-Detection-Range
max.-Detection-Range
Benedikt Merk-31.12.2020
CRS::EPSG:32632 - WGS 84 / UTM zone 32N - Projected
Detection Probability
0.0
0.2
0.4
0.6
0.8
1.0
0
200
400
600
Distance [m]


R2.2

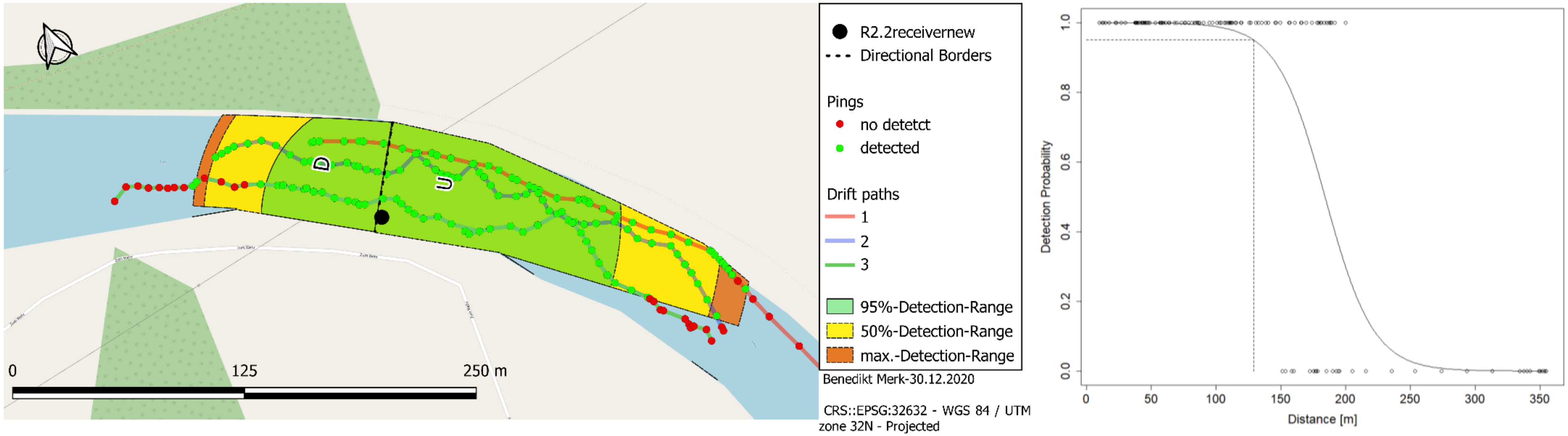

D
U
0
125
250 m
R2.2receivernew
Directional Borders
Pings
no detetct
detected
Drift paths
1
2
3
95%-Detection-Range
50%-Detection-Range
max.-Detection-Range
Benedikt Merk-30.12.2020
CRS::EPSG:32632 - WGS 84 / UTM zone 32N - Projected
Detection Probability
0.0
0.2
0.4
0.6
0.8
1.0
0
50
100
150
200
250
300
350
Distance [m]

## R1.1

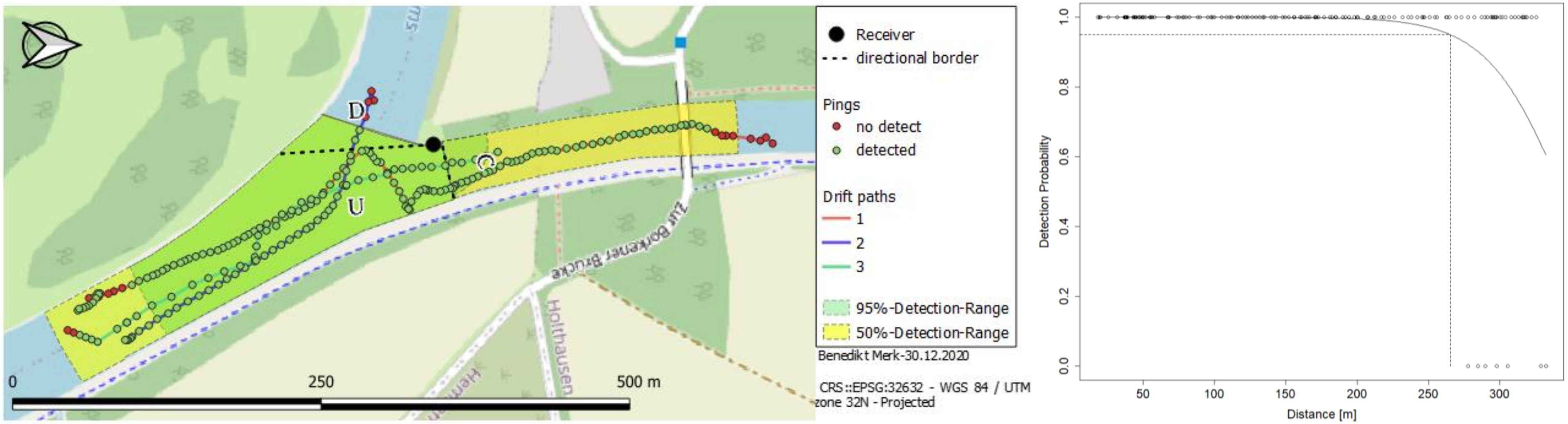
Receiver
directional border
Pings
no detect
detected
Drift paths
1
2
3
95%-Detection-Range
50%-Detection-Range
Benedikt Merk-30.12.2020
CRS::EPSG:32632 - WGS 84 / UTM zone 32N - Projected
D
C
U
0
250
500 m
Zur Borkener Brücke
Holthausen
Detection Probability
Distance [m]
0.0
0.2
0.4
0.6
0.8
1.0
50
100
150
200
250
300

## R1.3

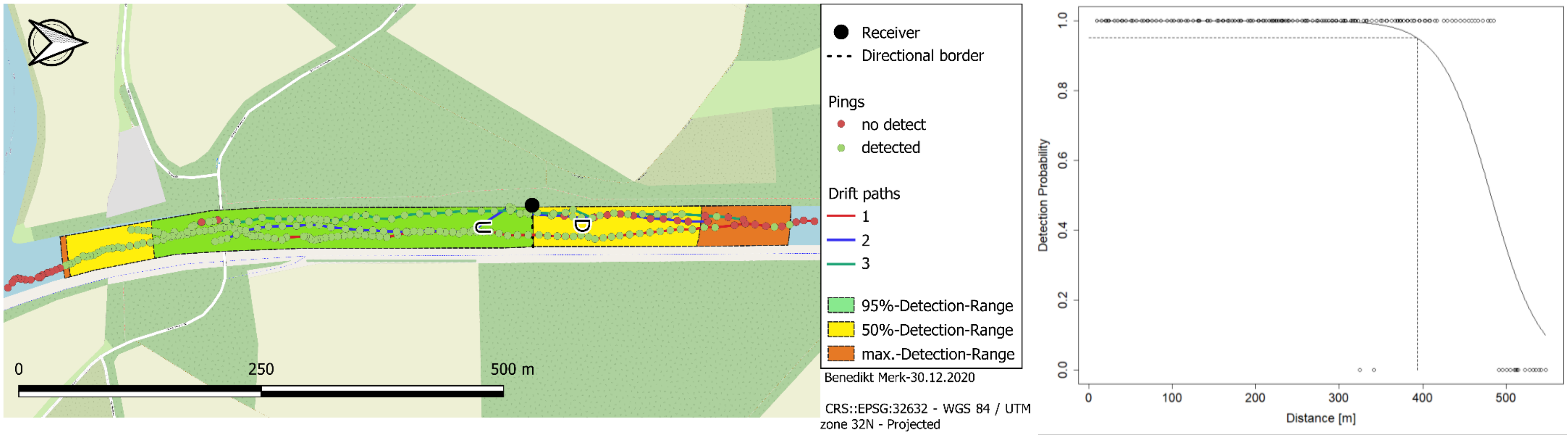
Receiver
Directional border
Pings
no detect
detected
Drift paths
1
2
3
95%-Detection-Range
50%-Detection-Range
max.-Detection-Range
Benedikt Merk-30.12.2020
CRS::EPSG:32632 - WGS 84 / UTM zone 32N - Projected
U
D
0
250
500 m
Detection Probability
Distance [m]
0.0
0.2
0.4
0.6
0.8
1.0
0
100
200
300
400
500

*Figure 4: Representation of the calculated detection by a map (left) and a graph (right) for each tested receiver. In the map includes the detection radii (green- 95%-detection-range, yellow-50%-detection-range, orange- absolute maximum detection range). The conducted drift routes are represented by a colored line (red- first drift, blue- second draft, green- third drift). Pings are categorized in detected (green) and non-detected (red). The receiver itself is indicated by a black dot. In case of directional separation, the border of the defined direction is marked by a dashed line. The top left arrow displays the north direction. The graph on the right is fitted with a binomial regression line. The 95%-Detection range is indicated by the intersect of the regression line to the dotted helper lines. In this figure just upstream drift are considered in the graphs. Other directions can be found in figure 5.*

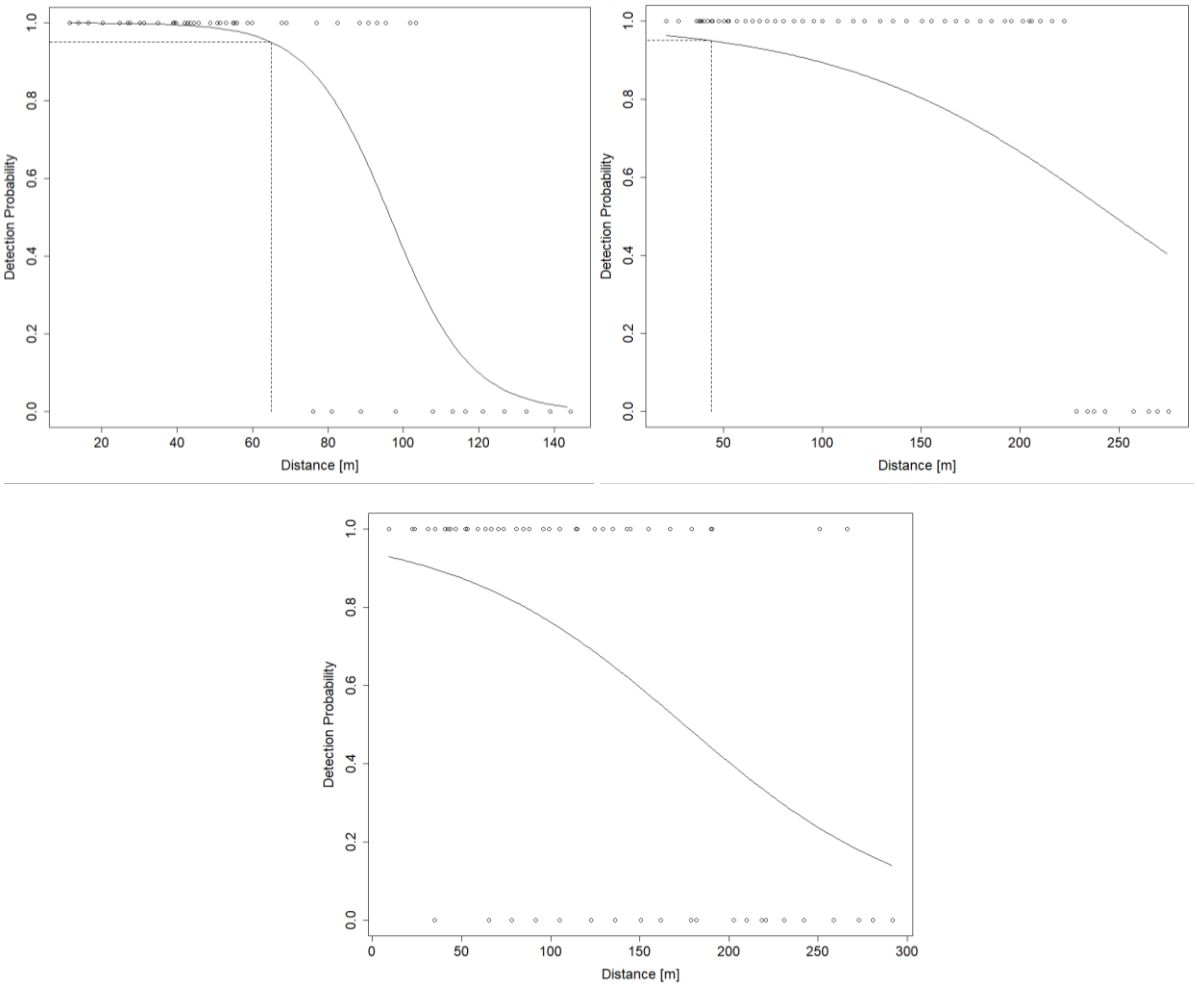


*Figure 5: Graphical representation of downstream/channel ranges of inland receivers. Intersection of regression lines and helper lines indicate 95%-detection-range. Top left: R2.2; Top right: R1.1; Bottom: R1.3*

The effect of the drift on the detectability was inconsistent among the receivers. No statistical influence of drift could be found at the channel direction of R1.1. An additive effect was validated for the receivers M1.1 as well as R2.2 and R1.3 in downstream direction, while the remaining tested receivers exhibited an interactive influence of drift and distance on the detectability (Appendix Table 5, Figure 9). This illustrates that the influence of distance on the detection probability depended on the drift.

#### 3.2. Tidal and Inland Receiver Comparison

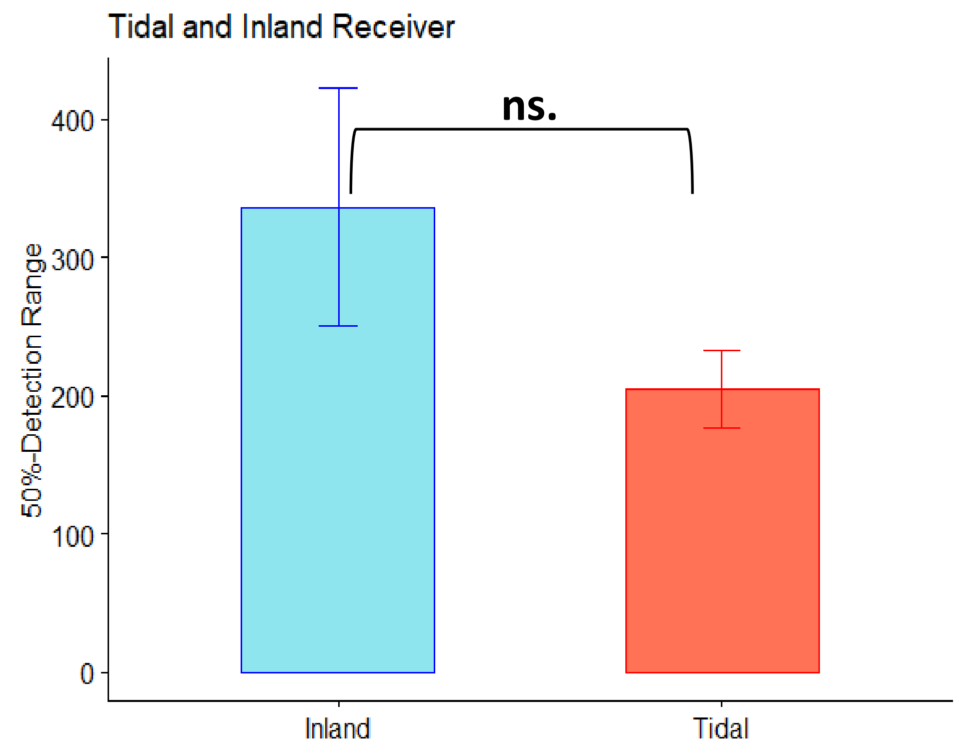


*Figure 6: Direct comparison of 50%-detection ranges of inland and tidal receivers*

The direct comparison of tidal and inland receivers revealed no significant difference ($p>0.05$, $\chi2_{(1,4)} =2.1343$). The mean inland detection radius was 336.47 ± 86.00 m exceeding that of the tidal receivers (204.28 ± 28.10 m).

### 4. Discussion

#### 4.1. Receiver Range-Test

*Table 3: Range diameter summary combined with eel velocity evaluation; + indicates a sufficient 95%-detection range to cover the distance an eel travels within the 60 second ping interval. * topographic irregularities (see Discussion). Distance travelled during 60 seconds: **-57m ***116 m*

| Receiver | 50%-Range-Diameter [m] | 95%-Range-Diameter [m] | Average Migration Speed ** | Maximum Migration Speed *** |
|---|---|---|---|---|
| D1.1 | 350 | 152 | +* | +* |
| D1.6 | 521 | 347 | + | + |
| M1.1 | 355 | NA | - | - |
| R2.2 | 281 | 194 | +* | +* |
| R1.1 | 591 | 309 | + | + |
| R1.3 | 656 | 394 | + | + |

The largest 95%- and 50%-detection diameter was observed at the inland receiver R1.3, with 394 and 656 m respectively. The absolute maximum detection distance of any receiver was M1.1 in the tidal area with 534 m under meteorological calm conditions (see Appendix Table 4). This maximum distance was slightly lower than 605 m measured by Béguer-Pon et al. (2018a) with a similar setup. However, this receiver exhibited no modelled 95%-Detection range. Compared to this study, the 50%-detectability range for V9 tags in the Gulf of St. Lawrence was between 250 and 300 m, thus slightly increased compared to this study. However, different receiver models (Vemco, VR2W and VR4) were deployed (Béguer-Pon et al. 2018a). A stationary range test study by Reubens et al. (2019) with VR2AR receivers at the Belgian long-term telemetry network revealed slightly lower ranges for receivers in tidal zones (max. range at 400 m and a sharp detection drop-off from 70% to 0% between 200 and 350 m). Ultimately, the 95%-detection diameter in this study exceeded the average and maximum migration distance a silver female eel can travel within 60 seconds for all tested receivers except M1.1. The calculated ranges can be transferred to other receivers within the BALANCE-project with similar conditions such as attachment method and surroundings.

The detection range of the autonomous receiver D1.1 in the tidal Dollart zone was insufficient to cover the distance to the shore line. While in river direction, the range largely overlaps with the next Dollart receiver D1.2, which is expected to have a comparable range as D1.6 (Figure7). Therefore, a relocation of this receiver closer to the shore line is recommended, to minimize chances of eels escaping in shallow waters at the shoreline outside the detection range of the receiver, as implied by the black arrow. It needs to be assured that the outer margin of D1.1 and the overlap between D1.1 and D1.2 offers a minimum length of 116 m. This ensures the detectability of migrating eels along the western coast in the Dollart estuary.

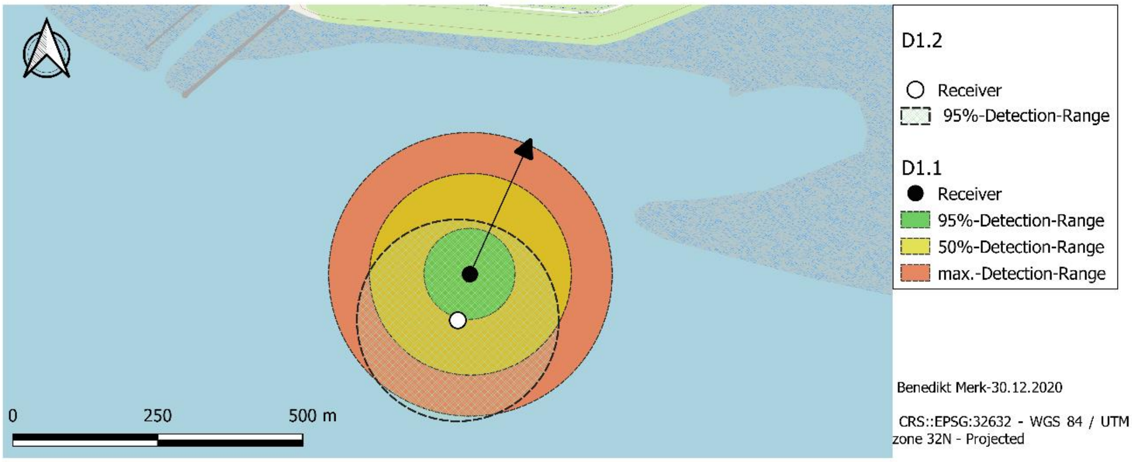


*Figure 7: Receiver D1.2 (white dot) and D1.1 (black dot) ranges; black arrow indicating a replacement a replacement in the target direction. Otherwise, legend entries are similar to Figure 4*

The tidal M1.1 receiver area is of special interest as many stow nets are located here, which are crucial for the eel data collection of the BALANCE-project. The first two drifts at M1.1 were conducted at unfavorable weather conditions with increased wind and wave heights, resulting in elevated background noises. Consequently, this lowers the number of detections in the first two drifts (Appendix, Table 4) (Kessel et al. 2014; Mathies et al. 2014; Reubens et al. 2019). The hard-shell boat type can possibly interfere with the acoustic signals. To avoid the influence of the boat, the tag should be placed at an increased depth in this case. Therefore, further investigation at different environmental conditions is recommended for this area for a more precise validation of the receivers' detection range.

Further, the detection range of receiver R2.2 needs to interpreted with caution. Firstly, the range was limited due to the river shoreline as the river Ems meanders. The calculated detection-ranges coincided with the distance of the receiver to the river curve. Secondly, the curve and shore line irregularities cast an acoustic shadow (Walsh et al. 2012), therefore the shortest possible distance an eel can travel through the receiver field with a 95%-detection probability is 130 m along the south-easter shoreline (Figure 4, map R2.2). However, this distance is sufficient for a reliable detection of a migrating eel.

Another irregularity occurred at receiver R1.3 in the "Dortmund-Ems"-channel in the downstream direction. The undetected pings in the downstream direction of drift 2 were located close to those of the detected pings of drift 3 (also indicated by the drift model R1.3 upstream see appendix). The only noted major difference was the presence of a vessel directly prior to the drift. While the motor noises showed no direct effect on the detection (compare pings of drift 2 close to R1.3 and retaining wall during the presence of the boat; Figure 4, map R1.3). The turbulences caused by the boat swirled up a lot of visible sediment which possibly intercept the pings.  As nautical traffic is a major subject at this location it is crucial to account for the effect of de-sedimentation for further studies. However, the

95%-detection-range was exceeding the maximum distance, which can be travelled by a migrating silver female per minute eel by far. Therefore, constrains to detectability from nautical traffic are unlikely.

In summary, the detection diameters of all receivers except M1.1 are sufficient to reliably detect female silver eels at various swimming velocities at least once as they are migrating through the 95%-detection field. Therefore, Hypothesis 1 can be validated for D1.1, D1.6, R2.2, R1.1 and R1.3. Minor adjustments to the placement of receiver D1.1 would minimize the chance of non-detects in the Dollart-Receiver-Chain along the western shore line.

#### 4.2. Inland and Tidal Receiver Comparison

Since M1.1 lacked a 95%-detection range, calculated 50%-detection radii were utilized for a more reliable comparison. Further restrictions of the comparison included the exclusion of directions the receiver's internal hydrophone was not facing towards, as this was the case for all inland receivers. The results illustrate a superior range of inland receivers of 336.47 ± 86.00 m (Figure 6). Nevertheless, the direct comparison of tidal and inland receivers revealed no significance between their 50%-detect-ranges. Consequently, hypothesis 2 needs to be rejected. However, this was most likely caused by the small number of replicates. Additionally, the range of the inland receiver R2.2 was limited by the river topography rather than documented environmental parameters (Figure 4, map and graph R2.2, Appendix Table 5). Therefore, a significant effect is expected with unobstructed receivers, since environmental parameters differ between these two river zones. Additionally, the GLMM (Generalized mixed effects model) exhibited a significant influence of distance on detection probability in dependence of tidal influence (Appendix Figure 9, Text 1). This reinforces the expectation.

This unconformity of detection ranges between inland and tidal receivers is assumed to be caused by environmental factors such as tidal currents and turbidity. A higher concentration of sediments in the water inhibits the sound transmission (Reubens et al. 2019). This was particularly evident after the de-sedimentation caused by the vessel during drift 2 at R1.3. Moreover, the buoy chain and the receiver tilted in the currents direction additionally decreasing the receiver's range (Reubens et al. 2019). Further, the noise of tidal currents between high and low tides overshadowed the pings of the tags. Thus, decreasing the effective detection range of the receivers (Mathies et al. 2014; Reubens et al. 2019). This was reinforced when considering the different drifts at D1.1 and M1.1 and their respective ranges. The third drift in both cases, conducted on tide with stagnating water flow, exhibited an enlarged range compared to Drift 1 and 2 at ebb/flood tide (Appendix Table 4). Consequently, a tidal effect through changing current velocities, tilting the receiver and increasing turbidity (Reubens et al.

2019) was expected to be a major influencing factor in the Ems estuary. Similarly, this was also the main effect in previous studies (Mathies et al. 2014; Reubens et al. 2019).

Additionally, the swell and winds could have been major noise sources especially in surface waters ( Kessel et al. 2014; Stocks et al. 2014). These environmental parameters were likely further influencing factors in the tidal zone. Moreover, the receivers in the tidal zone were attached to buoy chains. These were also sources of potential background noise close to the receiver, for example when the chain links collide. Particularly, this effect is present in combination with intensive wave action. Further, the increased salinity in tidal zones imply increased signal absorption (Crossin et al. 2017).

In contrast, inland areas were generally less influenced by tidal effects. Many artificial structures, like weirs and locks, decreased the visible current velocity drastically in the Ems river. Wave swell and wind were minor compared to the tidal conditions during this study. Further, the inland localities were typically expected to be less affected by coastal storms. Generally, the environmental conditions in the tidal independent zone favored higher acoustic ranges of the telemetry network. However, algal blooms and vegetation growth during spring and summer are known factors to inhibit signal transmission. In addition, seasonal changes in range due to stratification were also observed (Kessel et al. 2014; Mathies et al. 2014). Therefore, a lower detection range is anticipated during this time. Contrary, to the finding of the mentioned literature and indication by this study freshwater receivers can also have limited ranges due to river geometry and topographic obstructions (Walsh et al. 2012), as it was the case for one of the receivers (R2.2).

#### 4.3. Study Limitations

The range test tag was stabilized at a depth of 2 - 2.5 m, referring to pelagic conditions in most circumstances. Therefore, bathymetric irregularities castings acoustic shadows were not accounted in this experimental setup (Kessel et al. 2014). Consequently, eels travelling close to the bottom (Marohn et al. 2014) hiding between vegetation, stones and wood or simply lying on sediment might not be detected (Espinoza et al. 2011).

According to the drift models, range tests 1 and 2 at D1.1 conducted at similar environmental conditions resulted in different regression curves. Thus, hinting towards an influence of the perpendicular distance of drift path to receiver. The slope of the binomial graph decreases with increasing perpendicular distance of the drift path to the receiver, since more distance is covered within the transition zone of detects to/from non-detects (compare drift 1 to drift 2). However, the drifts of D1.6 contradict this argument. Therefore, to minimize those irregularities and allow for a superior quantification of the effect of different environmental parameters a clearly defined drift path is advised, especially in the more open Dollart region.

Due to increased current velocities during tidal changes the drift time and consequently the number of emitted pings was reduced for tidally influenced receivers as compared to those located inlands. Consequently, for increased current velocities the distance between pings is enlarged. Therefore, the 95%- and 50%-detections ranges are more inaccurate at shifting tides.

Since environmental parameters were not in focus of this study the respective effect sizes were inequal for all receivers. In particular, the conditions during the range tests of central-Dollart-receiver D1.6 were very similar and calm. Therefore, under unfavorable conditions a shrinkage of the effective detection diameter is expected. Additionally, with a higher quantity of drifts per receiver a more representative illustration of the environmental conditions can be depicted. However, extreme meteorological conditions cannot be covered with boat drifts due to safety issues (Kessel et al. 2014). Further, the represented receiver ranges are visualized as circles (Mathies et al. 2014). This could not be experimentally proven due to the limited drift number and restricted current directions.

A standard binomial GLM has a high model accuracy at center (50%)-values. Contrary, the regression line for upper and lower values (e.g., the 95%-range) is more imprecise (see Figure 5, R1.1) with an expected range of 170 m, contrary: a modeled 95%-detection-range of 43 m). This effect is more pronounced when artificial iterations are added (Table 5). Subsequently, this reasons for the missing 95%-Detection-Range of M1.1. Better model approaches for the 95%-Detection-Range could minimize these inaccuracies.

#### 4.4. Outlook

Nevertheless, this study can provide the basis for a long-term study on BALANCE-receiver network, as the results of this study indicate a strong dependence of the receiver range on environmental parameters. By quantifying those parameters, accurate predictions about the ranges at certain conditions can be calculated. Consequently, this results in a reliable foundation for interpreting behavior and quantity of migrating silver eels in the Ems river.

#### 4.5. Conclusion

Testing telemetry networks is crucial for quality control in telemetric studies and also the interpretation of biological linked data, such as migrating silver eels. An ideal telemetry network ensures that a passing tagged organism can be detected at least once during the tags ping interval. This study validated several receiver ranges and demonstrated that all tested, except M1.1, receivers exhibit a sufficient range to detect migrating silver eels. Nevertheless, further validations for the detection range of M1.1 and a relocation of D1.1 are recommended. The comparison of tidal and inland receivers illustrated no clear results. Therefore, future long-term range tests need to assess the impact of environmental influences in inland and tidal zones of the Ems river (Kessel et al. 2014; Mathies et al. 2014; Crossin et al. 2017; Béguer-Pon et al. 2018b). This would allow for more accurate depictions of the receiver ranges and provides a solid basis for interpreting biological data for future telemetry studies, such as the BALANCE-project.

### 6. Additional Material

*Table 4: Summary of conducted drift times and conditions during the range test*

| Receiver | Drift | Date | Drift time from… | to … | Drift time [hh:mm] | No of pings | No of detects | Detected pings [%] | Tidal conditions | Wind conditions | Wave conditions |
|---|---|---|---|---|---|---|---|---|---|---|---|
| D1.1 | 1 | 13.10.2020 | 11:24:58 | 11:37:48 | 00:12 | 30 | 10 | 33.3 | ebb tide | ++ | ++ |
| | 2 | 13.10.2020 | 12:08:41 | 12:21:49 | 00:13 | 28 | 6 | 21.4 | ebb tide | ++ | + |
| | 3 | 13.10.2020 | 15:58:23 | 16:13:03 | 00:14 | 30 | 14 | 46.7 | low tide | + | ++ |
| D1.6 | 1 | 14.10.2020 | 10:54:12 | 11:18:37 | 00:24 | 63 | 13 | 20.6 | ebb tide | o | o |
| | 2 | 14.10.2020 | 11:47:34 | 12:06:06 | 00:18 | 46 | 16 | 34.8 | ebb tide | + | + |
| | 3 | 14.10.2020 | 12:24:15 | 12:38:55 | 00:14 | 39 | 21 | 53.8 | ebb tide | + | + |
| M1.1 | 1 | 16.09.2020 | 16:27:44 | 16:52:49 | 00:25 | 59 | 9 | 15.3 | flood tide | ++ | ++ |
| | 2 | 23.09.2020 | 13:32:11 | 13:51:06 | 00:18 | 48 | 5 | 10.4 | flood tide | + | + |
| | 3 | 07.10.2020 | 15:00:20 | 15:32:45 | 00:32 | 75 | 45 | 60.0 | high tide | + | + |
| R2.2 | 1 | 12.10.2020 | 11:27:19 | 11:50:29 | 00:23 | 60 | 40 | 66.7 | - | o | o |
| | 2 | 03.11.2020 | 11:29:11 | 11:51:11 | 00:22 | 57 | 55 | 96.5 | - | + | o |
| | 3 | 03.11.2020 | 12:07:01 | 12:32:30 | 00:25 | 63 | 40 | 63.5 | - | + | o |
| R1.1 | 1 | 12.10.2020 | 16:15:30 | 17:06:27 | 00:50 | 127 | 114 | 89.7 | - | o | o |
| | 2 | 28.10.2020 | 17:04:33 | 17:18:34 | 00:14 | 40 | 35 | 87.5 | - | o | o |
| | 3 | 03.11.2020 | 14:30:37 | 14:46:50 | 00:16 | 43 | 41 | 95.3 | - | ++ | + |
| R1.3 | 1 | 12.10.2020 | 16:42:54 | 17:34:15 | 00:51 | 119 | 96 | 80.7 | - | o | o |
| | 2 | 28.10.2020 | 17:31:19 | 17:52:32 | 00:21 | 44 | 34 | 77.3 | - | o | o |
| | 3 | 03.11.2020 | 15:13:51 | 15:37:47 | 00:23 | 60 | 55 | 91.7 | - | ++ | + |

*Table 5: Statistical evaluation of all conducted models regarding the receiver test*

| Target receiver | Direction | Model type | MAM | artificial iterations added | p-Value | Distribution |
|---|---|---|---|---|---|---|
| D1.1 | - | General | Detects~Distance | no | 1.607e-15 *** | $\chi 2_{(1,86)}$=63.496 |
| | - | Drifts | Detects~Distance*Drifts | no | 0.004347 ** | $F_{(2,82)}$=10.877 |
| D1.6 | - | General | Detects~Distance | no | 2.644e-09 *** | $F_{(2,146)}$=35.43 |
| | - | Drifts | Detects~Distance*Drifts | no | 0.03176 * | $\chi 2_{(2,142)}$=6.899 |
| M1.1 | - | General | Detects~Distance | no | 8.174e-12 *** | $\chi 2_{(1,180)}$=46.724 |
| | - | Drifts | Detects~Distance+Drifts | yes | Distance: < 2.2e-16 ***, Drift: < 2.2e-16 *** | Distance: $F_{(1,178)}$=135.55, Drift: $F_{(2,178)}$=137.70 |
| R2.2 | Upstream | General | Detects~Distance | no | < 2.2e-16 *** | $F_{(1,127)}$=253.17 |
| | | Drifts | Detects~Distance*Drifts | yes | 0.007368 ** | $F_{(2,123)}$=9.821 |
| | Downstream | General | Detects~Distance | no | < 2.2e-16 *** | $F_{(1,49)}$=74.573 |
| | | Drifts | Detects~Distance+Drifts | no | Distance: < 2.2e-16 ***, Drift: 1.893e-06 *** | Distance: $F_{(1,47)}$=75.473, Drift: $F_{(2,47)}$=26.355 |
| R1.1 | Upstream | General | Detects~Distance | no | 3.223e-14 *** | $F_{(1,147)}$=57.593 |
| | | Drifts | Detects~Distance*Drifts | no | 6.831e-10 *** | $F_{(2,143)}$=42.209 |
| | Channel | General | Detects~Distance | yes | 4.739e-13 *** | $F_{(2,53)}$=52.31 |
| | | Drifts | Detects~Distance+Drifts | yes | >0.05 | - |
| R1.3 | Upstream | General | Detects~Distance | no | < 2.2e-16 *** | $\chi 2_{(1,162)}$=67.582 |
| | | Drifts | Detects~Distance*Drifts | yes | 0.0419 * | $\chi 2_{(2,158)}$=6.3449 |
| | Downstream | General | Detects~Distance | no | 0.0001018 *** | $\chi 2_{(1,57)}$=15.103 |
| | | Drifts | Detects~Distance+Drifts | no | Distance: 9.430e-10 ***, Drift: 2.671e-08 *** | Distance: $\chi 2_{(1,55)}$=37.439, Drift: $\chi 2_{(2,55)}$=34.876 |

D1.1

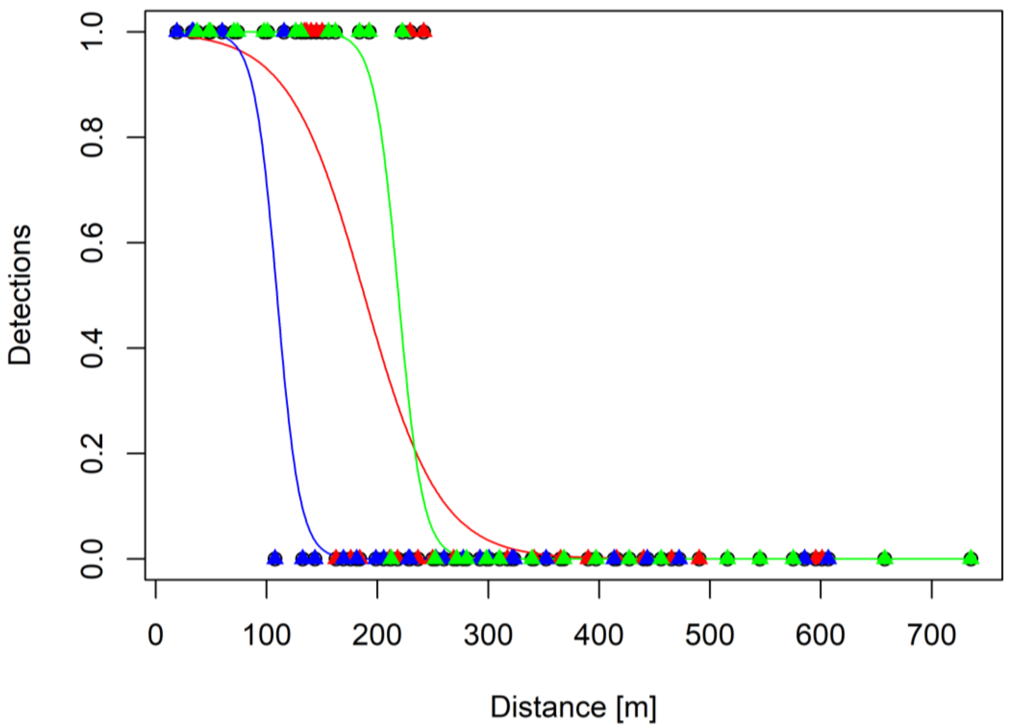
Detections
0.0
0.2
0.4
0.6
0.8
1.0
0
100
200
300
400
500
600
700
Distance [m]

D1.6

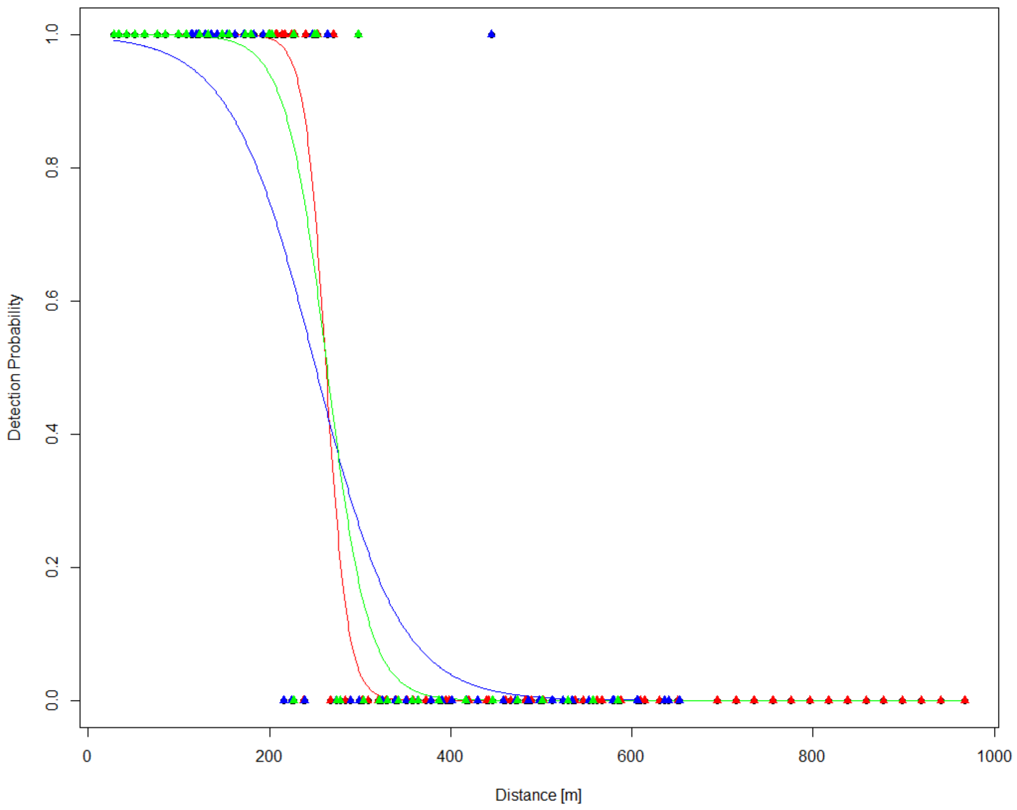
Detection Probability
0.0
0.2
0.4
0.6
0.8
1.0
0
200
400
600
800
1000
Distance [m]

M1.1

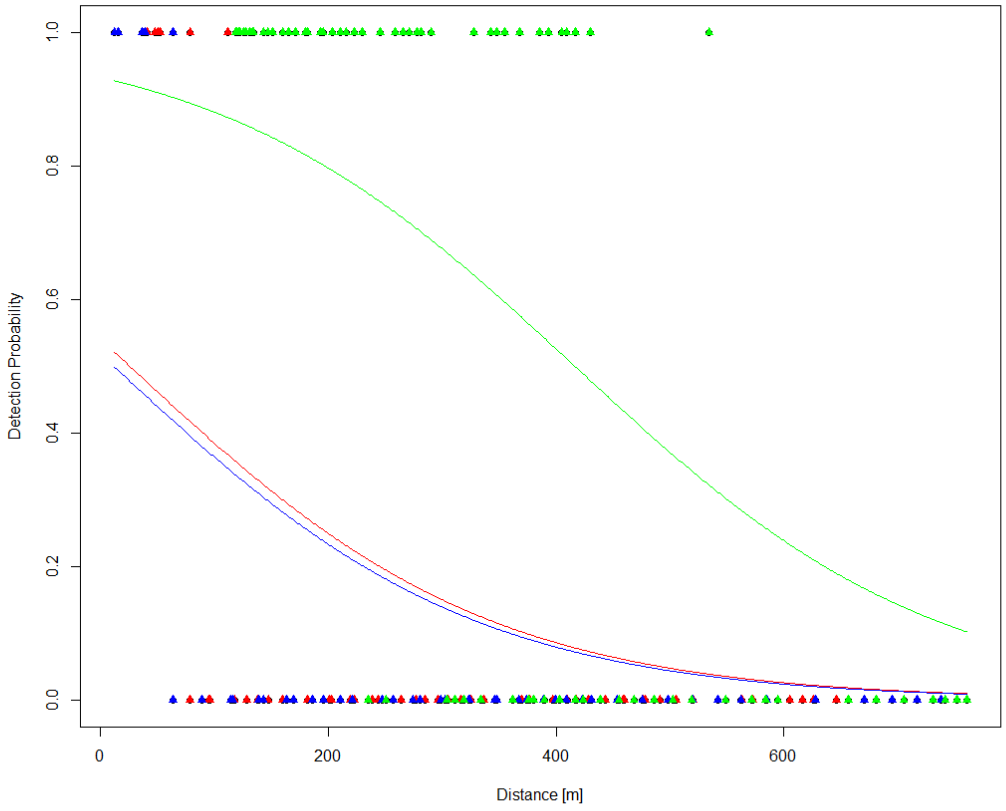


R2.2

Upstream

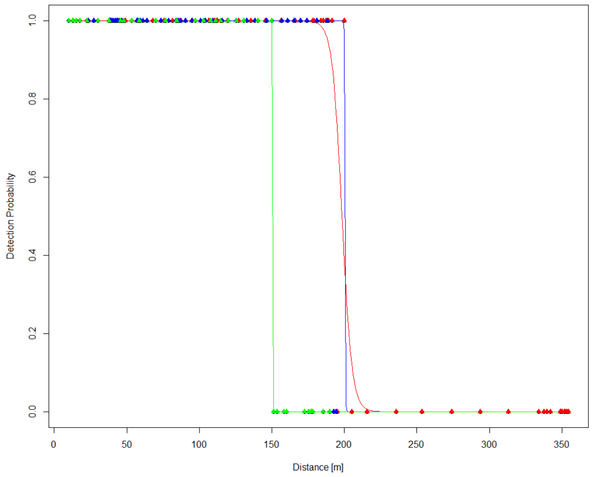


Downstream

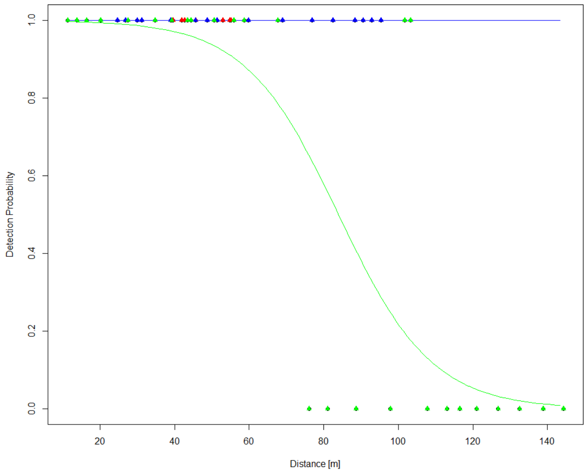

R1.1

Upstream

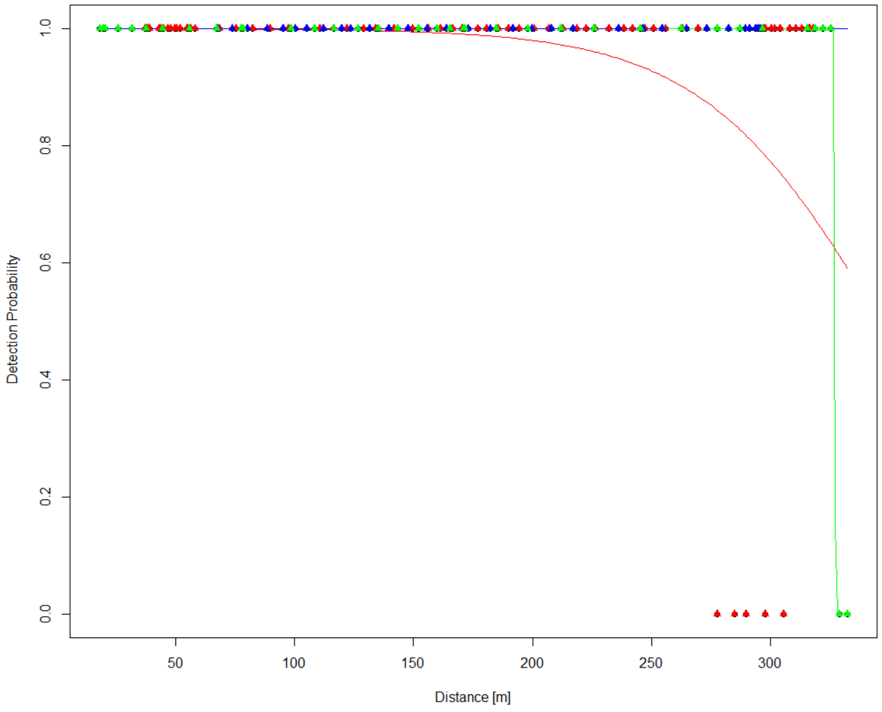


R1.3

Upstream

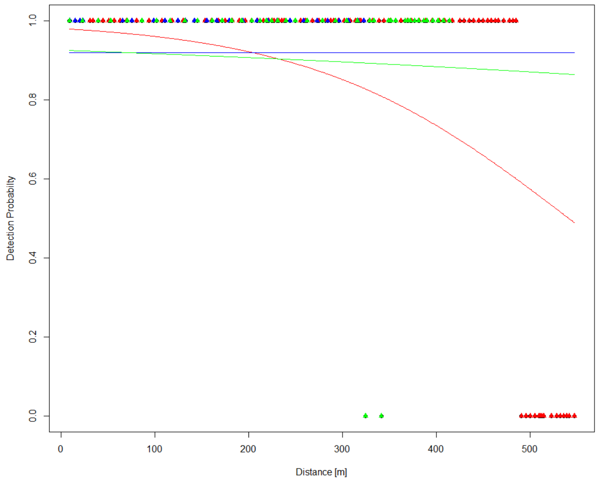


Downstream

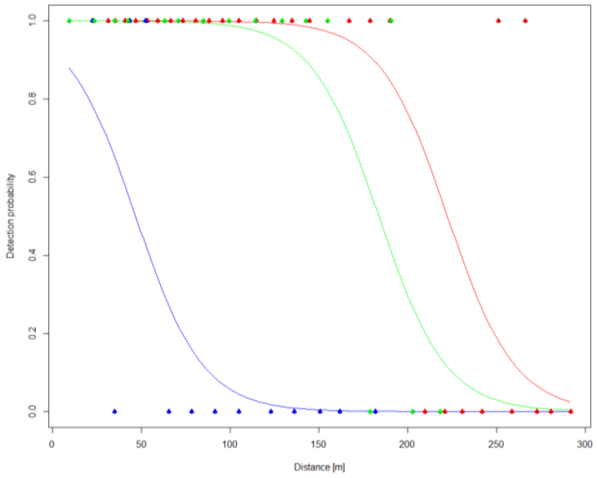


*Figure 8: Graphical representation of all drift models for all receivers. The drift colors are as followed: red- first drift; blue- second drift; green- third drift*

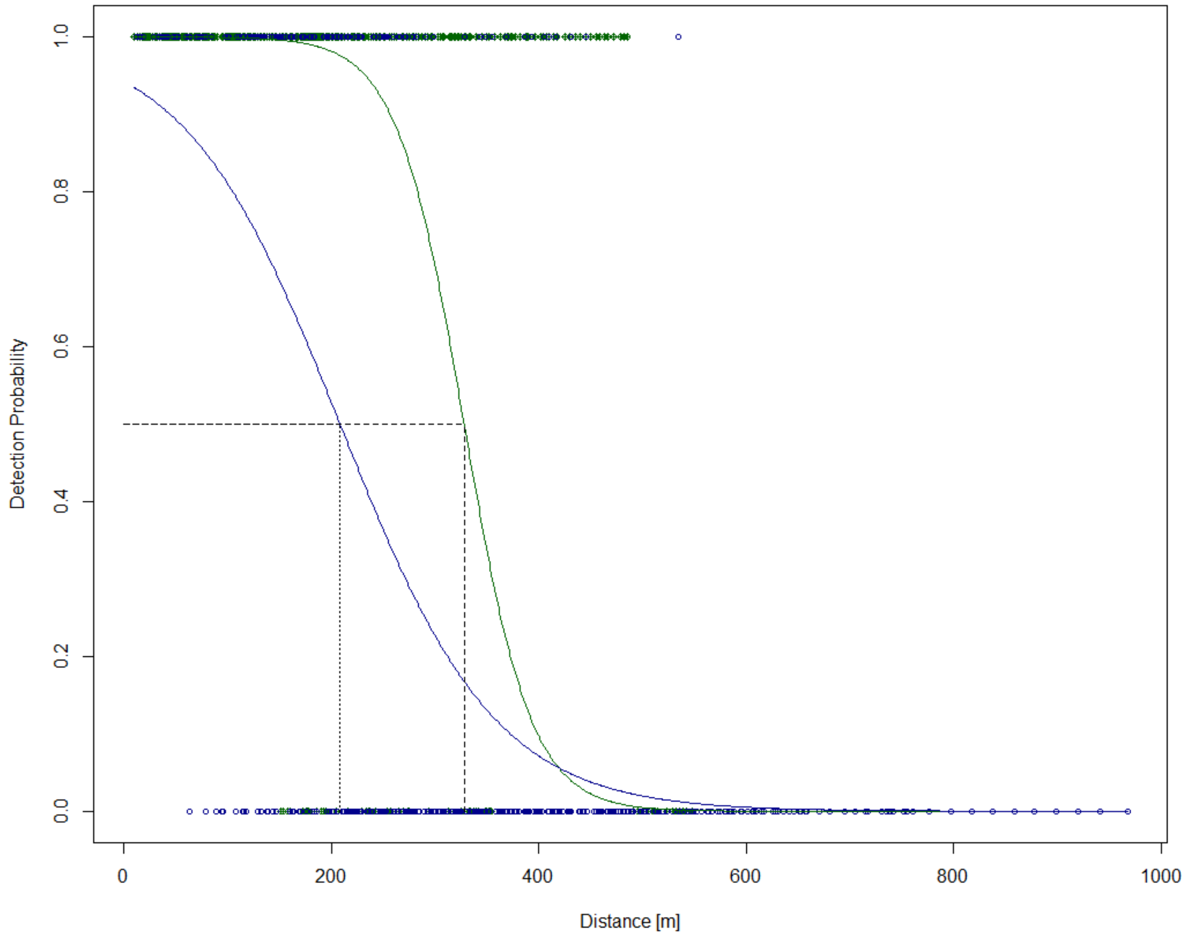


*Figure 9: Graphical representation of mixed model approach of the comparison of tidal (dark blue) and inland (dark green) receivers. The intersection of regression lines with dashed and dotted helper lines marks the 50%-detection range*

*Text 1:*

The generalized linear mixed effect model (GLMM) with binomial distribution revealed a significance of the interaction between range and tidal influence on the 50% detection-range (p= 0.0001575 ***, $\chi 2_{(1,862)}$ = 14.2799). Receiver are treated as random effect to weigh all receivers equally. The range calculation mostly coincides with the simple GLM with an inland range of 327.88 m and a tidal range of 208.37 m.

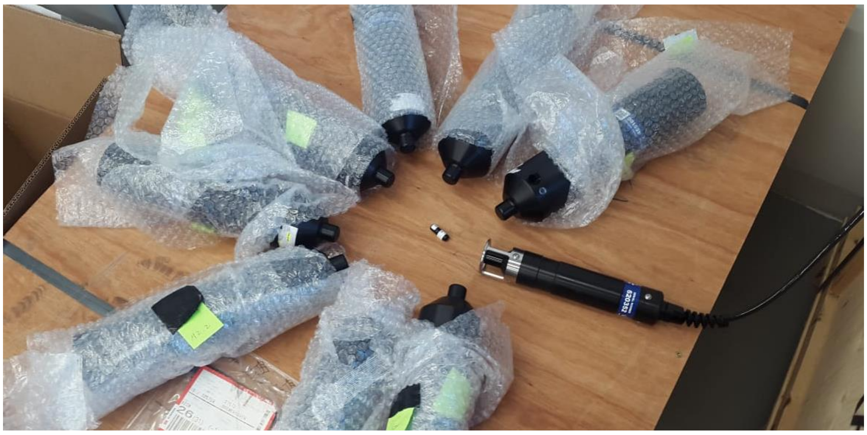

*Figure 10: Preliminary testing of VR2Tx receivers surrounding a V9 tag, tested by the VHTx-69kHz transponding hydrophone*
